# Distributed elasticity: a high-reward, moderate-risk strategy for efficient control modulation in insect flight

**DOI:** 10.64898/2026.03.23.713675

**Authors:** Liang Wang, Congxiao Zhang, Najmeh Asadimoghaddam, Arion Pons

## Abstract

The environments inhabited by flying insects demand a balance between flight efficiency and flight manoeuvrability. In structural oscillators such as the insect indirect flight motor, efficiency (arising from resonance) and manoeuvrability (arising from kinematic modulation) are typically *quid pro quo*, with modulation incurring penalties to efficiency. Band-type resonance is a phenomenon that offers, in theory, a strategy to lessen these penalties via careful navigation through a band of efficient kinematic states. However, identifying this band is challenging: no methods exist to identify the complete band in realistic motor models, involving elasticity distributed across thorax and wing. Nor are the effects of elasticity distribution on the band known. In this work, we address both open topics. We present a suite of numerical methods for identifying the complete resonance band in general systems. Applying them to models of the insect flight motor with distributed elasticity—thoracic and wing flexion—reveals that distributed elasticity is moderate-risk but high-reward morphological feature. Well-tuned distributions expand the resonance band over fourfold whereas poorly-tuned distributions completely extinguish the resonance band. These results indicate that distributing elasticity across the insect flight motor can have adaptive value, and motivate broader work identifying distributions across species.

## 1 Introduction

Insect flight motors are extraordinary natural vibrational propulsion systems. In a process that varies significantly across species, sets of muscles and exoskeletal mechanisms interact to generate finely-tuned wingbeat oscillation, and thereby, propulsive forces [1]. The efficiency and performance of these motors enable insects to carry out extreme feats of flight range, endurance, and manoeuvrability—single flights spanning hours to days and tens to thousands of kilometres [2, 3]; as well as complete body re-orientations on millisecond timescales [4, 5]. That insects can achieve this level of performance is a remarkable feature of flight motor operation—far surpassing the performance of state-of-the-art artificial micro-air-vehicles (MAVs), as illustrated in Fig. 1A. Many insect species are thought to utilise the integrative principle of structural resonance to improve flight efficiency: exoskeletal and muscular elasticity is thought to store and release wingbeat kinetic energy, leading to flight power reductions of up to 30% in certain species and conditions [6–8].

**Figure 1:**
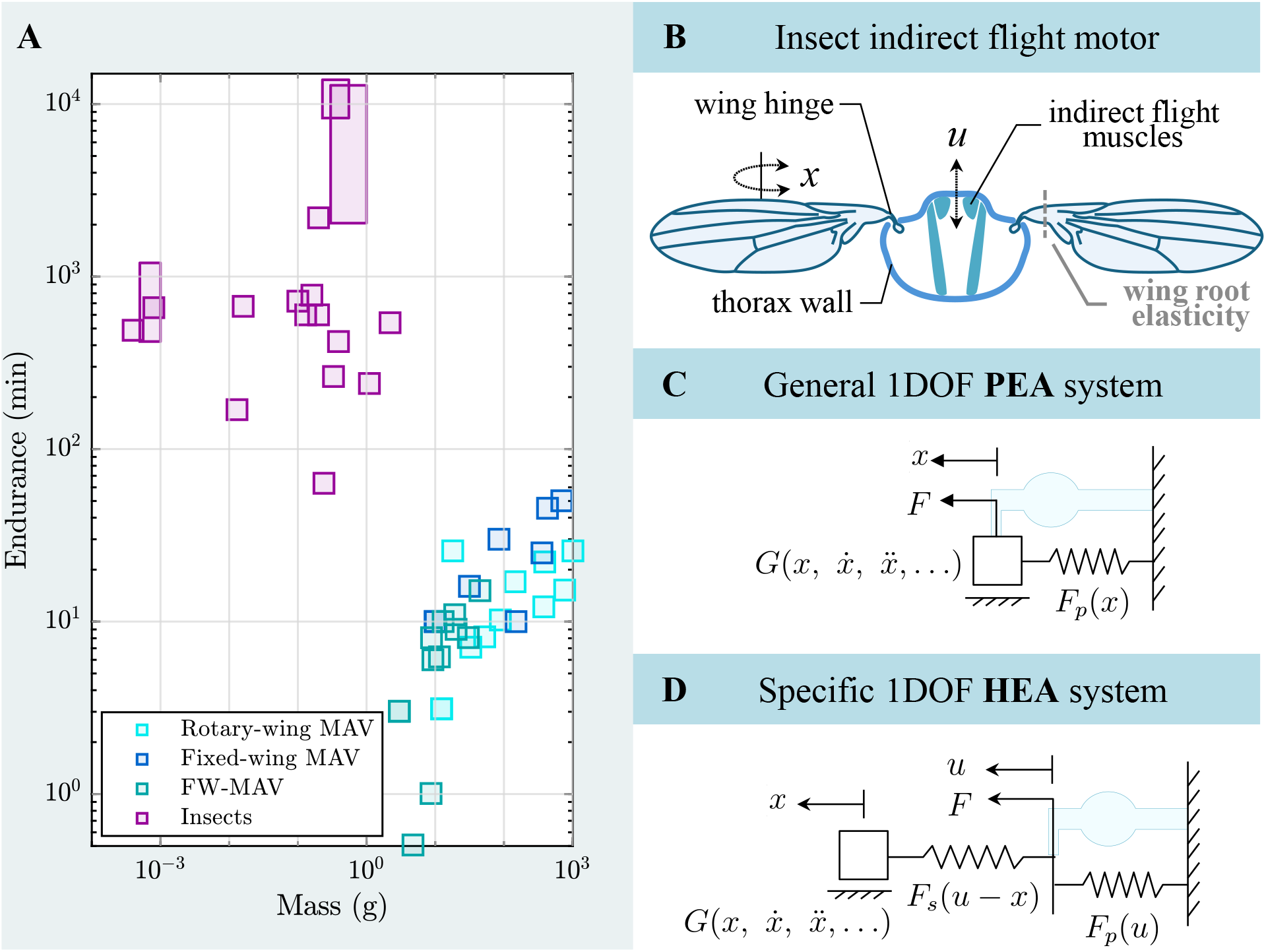
**A**.An illustrative comparison of flight endurances between insects and artificial MAVs. Insect endurances represent both intermittent and continuous flight, and are meta-analysed from [3, 33–40]. MAV endurances, as of 2015, are sourced from Floreano & Wood [41]. The significant superiority in insect endurances may reflect strategies for energy-efficient flight—motivating study of band-type resonance. **B**. Schematic of an insect indirect flight motor, illustrating the chordwise flexural line, *i*.*e*., wing root elasticity, described by Melis *et al*. [10] in drosophilids. **C**. General PEA system, as a candidate model of the flight motor. The parallel-elastic term *F*_*p*_ accounts for the combined elasticity of the primary flight muscles (illustrated in blue) and the thorax wall. **D**. A specific HEA system, accounting for chordwise flexion at the wing root via an additional series-elastic term *F*_*s*_: a distribution of elasticity between the thorax and wing root.

Elasticity plays an integral role in the generation of resonance, but its spatial distribution across the flight motor varies between species. In the classical conception of the insect indirect flight mechanism (as in Diptera), the elasticity of the primary flight muscles and that of the thoracic (pleural) wall are considered to lie in parallel: in mechanical terminology, a parallel-elastic actuation (PEA) system. However, recent studies have contemplated the additional presence of several forms of series elasticity, arising from thoracic local modes and/or flexion at the wing root [9–11]—features which can account for observations of a significant phase difference between muscular contraction and wing motion in some species. In addition, wing flexion—distributed across the wing, and acting along chordwise or spanwise axes—has long been recognised as a significant feature of certain species, notably bumblebees (*Bombus ignitus*) [12, 13] and hoverflies (*Eristalis tenax*) [14]. It is conjectured that these elasticities have a role in the storage and recovery of inertial energy—though, the significance of this role, and why distributions of elasticity vary between species, remain unclear [15].

As a yet further complication, while structural resonance can contribute to flight efficiency, this resonant efficiency appears inconsistent with the kinds of wingbeat control used to effect flight manoeuvrers, including wingbeat frequency modulation [16, 17], and symmetry/asymmetry modulation [18, 19]. For instance, a linear oscillator, deviation from simple-harmonic motion at a resonant frequency leads to losses in the classical transfer function magnitude associated with that resonant frequency [9]. Do insects simply accept these losses? Or are there other mechanisms at work? [9, 16]. A similar question arises in the design and control of insect-inspired flapping-wing micro-air-vehicles (FW-MAVs) operating on resonant principles [20–22]: does the use of resonance to improve the efficiency of these FW-MAVs preclude, or inhibit, classical FW-MAV flight control based on wingbeat modulation? [23]. How to modulate wingbeat kinematics while maintaining energy efficiency remains an open question in both insect and insect-inspired flight.

One phenomenon which sheds light on these questions is that of band-type resonance: the ability of both linear and nonlinear oscillators to achieve resonant levels of energy efficiency— a non-classical transfer function—not only at their discrete simple-harmonic resonant states, but within a continuous band of fundamental frequencies around these states [24, 25]. This invariant efficiency is achieved via a spectral shaping process, in which the input forcing waveforms are tuned away from simple harmonic to ensure that power flows only from the forcing actuator to the oscillator, and never in reverse: a condition of global, or energy resonance [26]. By operating in this band, insect and insect-inspired propulsion systems could modulate wingbeat kinematics with no penalty to mechanical efficiency. However, to date, our understanding of band-type resonance is limited to conservative analytical estimates for simple systems [24, 27] that do not reflect the complex distribution of elasticity present in the flight motors of many insect species. Classical transfer function analysis indicates that realistic distributions of motor elasticity can leave to an expansion in the acceptable wingbeat frequency modulation window [9]—motivating a hypothesis the energy resonance band might also expand. However, to date, no general method exists to compute the full resonance band even in linear oscillators, leaving this hypothesis untestable.

In this work, we overcome this methodological barrier and test this hypothesis. In §2, aggre-gating a range of experimental data, we identify a series of dynamical systems representing the indirect flight motor with distributed elasticity, focusing on the chordwise flexion at the wing root observed by Melis *et al*. [10] in drosophilids. In §3 we develop a suite of general numerical methods for mapping the set of band-type resonant states, based on particle swarm optimisation (PSO), numerical continuation, and the combination of these techniques. Applying these methods to our models, we obtain (§4) complete maps of their frequency-band resonant states: energy resonant states across wingbeat frequency under symmetric wing-beat kinematics. These maps yield several surprising insights. Among them, they reveal that band-type resonance is not always contiguous: in heavily-damped systems, isolated regions of energy resonant and near-energy resonant states can re-emerge at very low frequencies (down to 20% of the natural frequency). This demonstrates that the energetic penalties of wingbeat frequency modulation are not necessarily monotonic: larger modulations can occur lower energetic penalties than smaller ones. They reveal also that the boundaries (*i*.*e*., extremal wingbeat frequencies) of the contiguous resonance band are controlled by a simple muscle force timing relation—a prospective open-loop control strategy for obtaining efficient wingbeat frequency modulation in an insect or FW-MAV. And finally, they reveal that distributing elasticity across the flight motor is a moderate-risk but high-reward strategy for improving the efficiency of wingbeat frequency modulation. When the elasticity distribution—between thorax and wing root—is tuned optimally to the relative strength of damping ratio in the motor (the Weis-Fogh number), the resonance band expands massively: over fourfold with respect to the non-distributed case. However, when the distribution is mistuned, the resonance band can cease to exist entirely—potentially, rendering a flight motor based on muscular stretch-activation completely inoperable. Interpreting estimates of elasticity distribution in drosophilids, we find that they benefit from this distribution but are not tuned optimally. These results provide new insight into the adaptive value of distributed elasticity in the insect flight motor, as well as possible explanations for differences in elasticity distribution between species. There is a need for further comparative studies of these differences in distribution—as well as a possibility for new efficient control strategies for FW-MAVs.

## 2 Indirect flight motor models

### 2.1 Elastic actuation systems and elasticity distribution

A conventional initial model for an insect indirect flight motor, as illustrated in Fig. 1B-C, corresponds to parallel-elastic actuation (PEA). Here, the parallel elasticity is that of the pleural wall and of the primary flight muscles, and the wingstroke and muscles are rigidly linked, *i*.*e*., locked at zero phase lag. This phase lock is a characteristic that discriminates between insect species that are well-modelled by PEA and those that are not—though, observational data is sparse. Data by Iwamoto & Yagi [28] for bumblebees (*Bombus ignitus*) shows near-zero phase lag between muscle contraction and wingstroke motion, indicating PEA model applicability—though results by Cote *et al*. [29] in an unidentified *Bombus* sp. call this into question. Parallel elasticity is also a feature of several current FW-MAV platforms [23, 30, 31]. As per Fig. 1C, a general single-degree-of-freedom (1DOF), nonlinear, time-invariant, PEA system can be modelled:

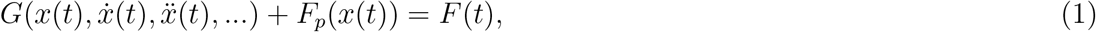

with time variable *t*, force input *F* (*t*), and wingstroke motion *x*(*t*).

The phase lags between muscle and wingstroke that are observed in other species of insects indicate, however, that PEA models are not applicable to all flight motors. Previously, Pons & Beatus [9] modelled a motor with a thoracic local mode, *i*.*e*., additional elasticity between the muscle and wing hinge, to account for these lags. However, recent observations by Melis *et al*. [10], point rather to a line of chordwise wing flexion near the wing root, *i*.*e*., additional elasticity between wing hinge and the bulk of the wing, as their origin—at least in drosophilids. In hawkmoths, *Agrius convolvuli*, significant phase lags are also observed [9, 32], with unclear structural origins. A general hybrid elastic actuation (HEA) system accounting for both classical parallel elasticity and the series elasticity of flexion near the wing root is illustrated in Fig. 1D, and may be described as:

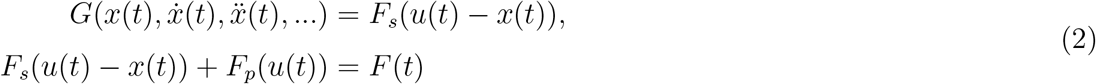

where *u*(*t*) is the muscular contraction and *x*(*t*) is the wingstroke motion—kinematic variables which are now no longer rigidly linked.

To describe specific forms of these general models, we define two functionals, *i*.*e*., functions of functions: *F* (*t*) = 𝒟{*x*(*t*)}, the actuator force in terms of output motion; and 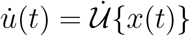, the actuator velocity in terms of output motion. The relevance of these two functionals is that, together, they determine the actuator mechanical power output, as 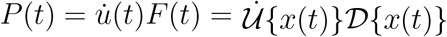. For PEA, 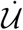 is the identity (*u* = *x*), but for HEA system it is more complex. To the construction of these functionals we now turn.

### 2.2 Linear and nonlinear model components

Three broad physical effects within the functionals of §2.1 should be described: (**i**) inertia, largely of the wings but with a small thoracic component; (**ii**) damping, largely arising from wing aerodynamic drag, but with a small thoracic structural component; and (**iii**) elasticity. In this work, we consider linear inertia and linear elasticity. There is evidence for weak elastic nonlinearities in the exoskeleta of honeybees (*Apis mellifera*) [42] as well as in the overall flight motor of drosophilids [6]. Such weak nonlinearities can be linearised with minimal impact on motor dynamics [43]. Turning to damping, we consider two models. The scaling of aerodynamic and, structural damping with wingbeat frequency is complex [44–46]: overall, this scaling is likely greater than linear but weaker than quadratic.

First, for validation and theoretical insight, we consider linear damping—leading to the canonical linear PEA oscillator:

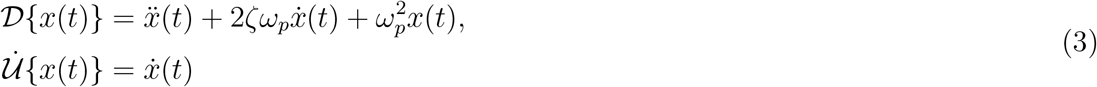

where *ζ* is the linear damping ratio, and *ω*_*p*_ is the motor natural frequency. We take *x*(*t*) as the normalised wing stroke angle: −1 ≤ *x* ≤ 1 for normative hovering flight. More realistic, however, is the case of quadratic damping:

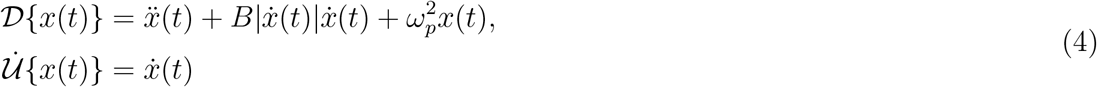

with quadratic damping ratio *B*. Taking again −1 ≤ *x* ≤ 1 for normative hovering flight, *B* may be approximately identified with the parameters 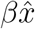 and *c* identified by Pons *et al*. [9, 4*3], and the parameter 1/N*_WF_ where *N*_WF_ is the Weis-Fogh number, as per Lynch *et al*. [46]. The caveat to this approximation is that these latter parameters assume varying levels of simple-harmonic behaviour within the motor—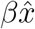, simple-harmonic *x*; *c* and *N*_WF_, simple-harmonic *x* and *F* —whereas we will specifically be considering multiharmonic waveforms. Nonetheless, these estimates indicate that we should expect 0.2 ≤ *B* ≤ 1.1 for the flight motors of various insect species [9]. In Appendix A.4, we estimate this parameter for several insect species.

Turning to the HEA system, Eq. (2), we consider only quadratic damping. With some manipulation, we may express the system functionals as:

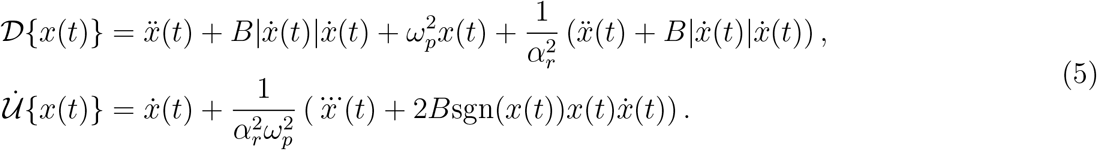

where *α*_*r*_ = *ω*_*s*_*/ω*_*p*_ describes the elasticity ratio between the the wing chordwise flexion (series-elastic, *ω*_*s*_) and thoracic/muscular (parallel-elastic, *ω*_*p*_) natural frequencies. The limit *α*_*r*_ → ∞ recovers the PEA system, Eq. (4). Note that this parameter is analogous, but not identical, to the parameter *α* identified by Pons & Beatus [9]: this latter *α* models thoracic local deformation, not wing flexion. In Appendix A.5, we estimate *α*_*r*_ for several insect species.

### 2.3 Actuator mechanical efficiency

For all of these motor systems, the actuator power output can be positive (*P >* 0), negative (*P <* 0), or zero at an instant. Positive power output refers to power flowing from the actuator to the system—increasing the total energy of the system. Negative power output refers to the reverse—power flowing from system back to actuator, reducing system total energy. Intuitively, reducing the level of negative power output increases the system efficiency, in a certain sense: it is inefficient to use the actuator to withdraw energy from the system, *cf*. [47]. In the best possible case, the actuator itself might imperfectly store this energy in an internal potential (*e*.*g*., in a regenerative actuator [47]); but in the worst case, the actuator’s own power source is consumed in order to withdraw this energy (*e*.*g*., as resistive power losses in an electromagnetic actuator). This intuition can be formalised [24, 25]: in a wide range of systems, 𝒟 (·), and kinematics, *x*(*t*) or equivalently *x*(*τ*), the state of non-negative power (*P* (*t*) ≥ 0, ∀*t*) represents optimality in absolute mechanical power consumption 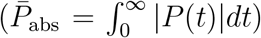 with respect to elasticity, *F*_*p*_(*x*). That is, it is not possible to alter the system elasticity, *F*_*p*_(*x*), such that 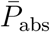 is reduced—and the same applies to a range of other power consumption metrics, covering the behaviour of a range of physical actuators [24]. This condition of optimality has been termed the global resonance [26] or energy resonance [43] condition. Defining the mechanical efficiency of the actuator, *η*, as the ratio between the net power 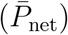 and the absolute power:

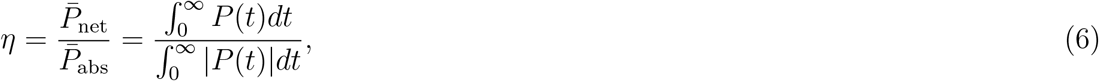

we see that energy resonance corresponds to *η* = 1.

### 2.4 Band-type resonant states

The set of periodic states, *x*(*t*) ∈ 𝒳, for which *η* = 1 are the *band-type resonant states*. This set includes the conventional energy resonant state, if one exists: the state outputting simple-harmonic motion, 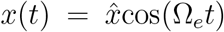. This frequency, Ω_*e*_, is the system’s energy-resonant frequency. For both linear and nonlinear PEA, Eqs. (3) and (4), the energy resonant frequency is the undamped natural frequency, Ω_*e*_ = *ω*_*p*_ [9]. However, *x*(*t*) ∈ 𝒳 includes also states outputting multi-harmonic waveforms with fundamental frequencies (Ω) away from Ω_*e*_: the frequency-band resonant states. In an insect flight motor or FW-MAV, these frequency-band states can enable energy-efficient high-authority control of wingbeat lift forces, via modulation of the wingbeat fundamental frequency, *cf*. [23]. In both cases, the frequency interval, Ω_min_ ≤ Ω ≤ Ω_max_, over which these states exist determines the authority of this mode of control. Determining this interval is difficult: it involves searching over all possible periodic waveforms at a given fundamental frequency (Ω) to either find a state with non-negative power, or confirm that no such state exists.

Previous analytical results [24] indicate that band-type resonant states for the linear PEA system exist over a frequency window of at least:

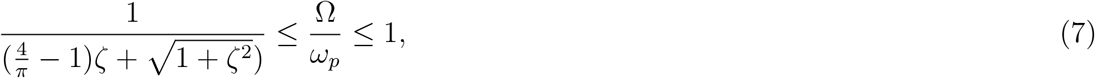

which, for damping ratios roughly commensurate with insect flight motors (0.11 ≤ *ζ* ≤ 0.64 [9, 43]), implies that the wingbeat frequency could conceivably drop to as low as 70% of the motor natural frequency before mechanical efficiency is lost. Eq. (7) is, however, a conservative result: the complete resonance band is anticipated to be larger, but existing techniques are insufficient to fully map it. In addition, the nonlinear PEA system and HEA system with distributed elasticity have not been characterised, and these systems appear less amenable to analytical approaches. For this reason a methodological advance is required, which we now provide.

## 3 Numerical methods for mapping band-type resonance

### 3.1 Waveform description

To define and solve the problem of mapping the set of band-type resonant states within the motor models of §2.2 we require a general parametric description of the waveform *x*(*τ*). Considering a general periodic oscillation with angular fundamental frequency Ω, we can define a nondimensional time *τ* = Ω*t*, which ranges from 0 to 2*π* over the period—a wingbeat cycle parameter. A natural candidate for the approximation of smooth periodic waveforms is a Fourier series. We consider a series of the form:

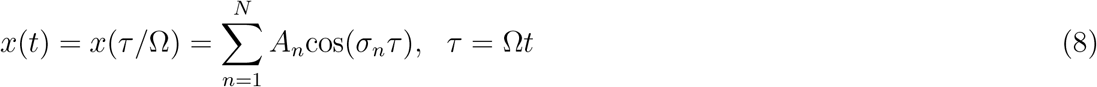

with odd harmonics, ***σ*** = [1, 3, 5, …, 2*N* + 1], where *N* is the number of harmonics. The coefficients **A** = [*A*_1_, *A*_2_, …, *A*_*N*_], and the frequency Ω defining *τ*, represent the parameter space of the waveform. As a cosine-only series, Eq. (8) already restricts the waveform parameterisation to waves which are even-symmetric about the nominal stroke extrema (*τ* = 0, *π*, 2*π*), *i*.*e*., to a wingbeat with symmetric upstroke and downstroke. The odd-harmonic series adds an additional restriction to waves which are odd-symmetric about the quarter-stroke points (*τ* = *π/*2, 3*π/*2), *i*.*e*., to wingbeats in which each quarter-stroke is identical, up to the appropriate change in sign. The Dickinson-type simplified wingstroke pattern [48], as used by Pons & Beatus [24] to derive Eq. (7), is one such odd-harmonic wave. Provided that the other wingbeat kinematic variables show the same quarter-stroke symmetry, the wing geometry is symmetric, and the wing aerodynamics are sufficiently low-order (neglecting, *e*.*g*., the influence of the body and effect of turbulence), odd-harmonic wingbeats will generate aerodynamic forces that are perfectly balanced around the midstroke point: with a net lift force but no other net aerodynamic forces on the insect or FW-MAV, considerably simplifying flight control.

The waveform description of Eq. (8) does, however, come with several caveats. First, the Fourier-series formulation limits the accuracy with which we can approximate non-smooth waveforms—including the Dickinson-type pattern. Second, if we consider several waves at differing fundamental frequencies Ω_*i*_, then we observe an aliasing effect to occur. An dense-harmonic wave at fundamental frequency Ω_1_ and with coefficients [*A*_1_, *A*_2_, …] is exactly identical, in the time (*t*) domain, to a sparser wave at one third the fundamental frequency, Ω_2_ = Ω_1_/3, with coefficients [0, *A*_1_, 0, *A*_2_…]; and to a wave at Ω_3_ = Ω_1_/5, with coefficients [0, 0, *A*_1_, 0, 0, *A*_2_…], and so on. Any wave can be recovered at certain lower nominal fundamental frequencies by setting the lower-frequency amplitudes to zero and using higher harmonics to alias back onto the original wave. To prevent this, and ensure that the fundamental frequency Ω is representative of the actual dominant period of the wingbeat, we will have to constrain its amplitude (*A*_1_) during problem formulation (§3.2).

### 3.2 Mapping problem formulation

With the waveform description of §3.1, we can formulate a problem for mapping band-type resonant states that is solvable via numerical methods. Given a selection of harmonics ***σ*** of length *N*, we seek pairs of frequencies, Ω and coefficient vectors **A** = [*A*_1_, …, *A*_*N*_] such that the system satisfies:

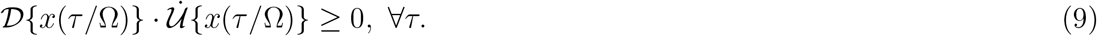

With the waveform parameterisation of Eq. (8), the derivatives of *x*(*t*) in 𝒟{·} and 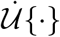 can be evaluated exactly, without discretisation, via:

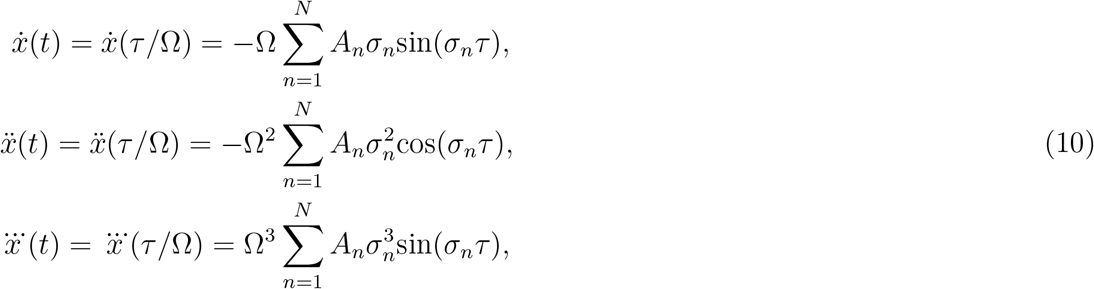

and so on. Based on existing characterisations of band-type resonance [24], we expect that the space of solutions, {Ω, **A**}, to Eq. (9) will be complex. For any given system, over Ω, we expect there to be regions within which there are multiple solutions: within which there is no solution; and (possibly) within which there is a unique solution. Insofar as the system dynamics 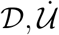 are dependent on parameters (*ζ, B, α*_*r*_), we seek the relationship between the properties of this solution region and these parameters—a map of the solution space.

### 3.3 Local constrained optimisation

We begin by formulating a method to solve Eq. (8) at only a single Ω = Ω_*i*_ based on local constrained optimisation. We discretise dimensionless time *τ* as ***τ*** = [0, 2*π/*(*L*− 1), 4*π/*(*L*− 1), …, 2*π*] with length *L*. For any selected **A**_*k*_ we may then evaluate the kinematic vectors 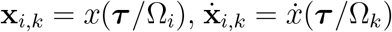, *etc*. (Eq. (8), (10)) and the corresponding vector of actuator power output (**p**_*k*_) via evaluation of the system functionals pointwise over the kinematic vectors (*i*.*e*., in loose notation, 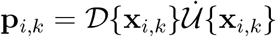. With the Fourier formulation providing time derivatives, this process is exact at the chosen time points ***τ***. We extract the minimum and maximum values of **p**_*i,k*_ and **x**_*i,k*_, and define a power ratio metric *γ*_*i,k*_ together with an amplitude metric *r*_*i,k*_:

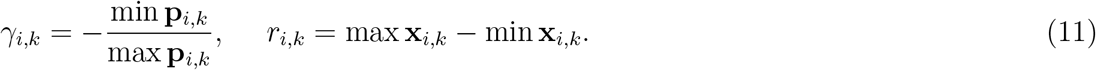

These metrics are now numerically-computable functions of Ω_*i*_ **A**_*k*_.

At the specified Ω = Ω_*i*_, we then use sequential quadratic programming (SQP) to minimise *γ*_*i,k*_ over **A**_*k*_, subject to the constraints of normalised wingstroke amplitude, 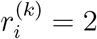, and the dominance of the fundamental frequency (*A*_1,*k*_)^2^ ≥ 1/4 (*i*.*e*., |*A*_1,*k*_| > 1/2, with normalized amplitude). If the resulting minimised *γ*_*i,k*_ is itself sufficiently close to zero (*γ*_*i,k*_ < *ϵ, ϵ* = 1 × 10^−6^ in our optimization) then we classify the optimised state as a band-type resonant state. In concise form, the full local optimisation problem can be described as:

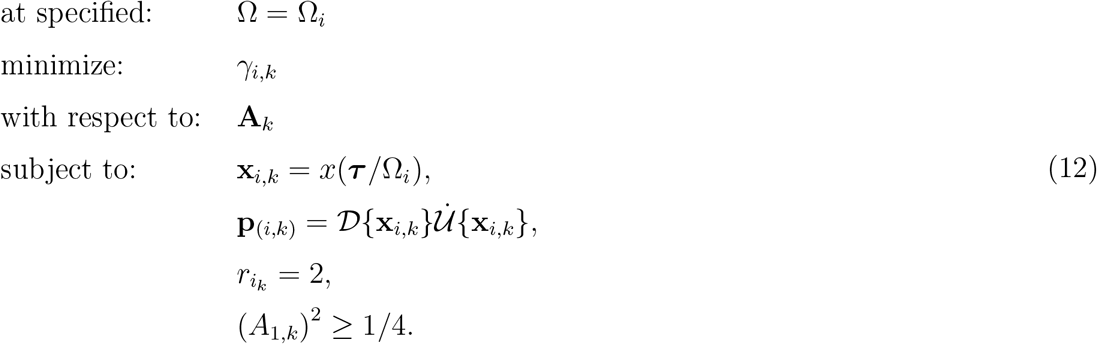

SQP is an iterative method, and so a suitable initial guess (**A**_*k*=0_) is key to solver performance— this leads us to the topic of how to generalise local optimisation to global mapping.

### 3.4 Natural continuation for mapping

Analytical results [24] indicate that band-type resonance has a certain structure. For the PEA systems we consider (Eq. (3)-(4)), there is a simple harmonic solution, **A**_0_ = [1, 0, …], at *ω*_0_ = *ω*_*p*_. In the case of the HEA system, the equivalent simple harmonic solution, *ω*_0_ (if it exists) can be found via a numerical parameter sweep over Ω. These simple-harmonic states provide an ideal start point for a mapping method based on numerical continuation. Starting from them, we increment Ω and use the previous solution as the initial guess for the local optimisation. An algorithm in pseudocode is shown in Appendix A.1. Starting from the simple-harmonic state (line 1), we separately conduct the upward search (line 2, (*ω*_0_, Ω_max_)) by consecutively increasing the frequency (line 16, ′+′) and the downward search (line 2, (Ω_min_, *ω*_0_)) by consecutively decreasing the frequency (line 16, ′−′). If the solution fails at any point, the preceding frequency is designated as the boundary frequency, Ω_*B*,up_ for the up and Ω_*B*,down_ for the down (lines 13), and the remaining area will not be further explored (lines 14). For this reason, this approach for mapping is highly time-efficient, but can only explore a contiguous solution region.

### 3.5 A particle swarm method for mapping

There is, however, no guarantee that the resonance band is contiguous. As a more general, but time-expensive, approach to global mapping, we apply a particle swarm method. We generate a large population (size *M*) of randomly distributed initial guesses (particles, **A**_*i,k,j*_, 1 ≤ *j* ≤ *M*) and apply local constrained optimisation to each particle. An algorithm in pseudocode is shown in Appendix A.2. The matrix **A**_**In**_ with *M* rows (line 3) stores the initial guesses for each particle. Unlike natural continuation, this particle swarm method approach explores the entire frequency range of interest, (Ω_min_, Ω_max_), and is capable of identifying non-contiguous regions. However, solutions may not be continuous in **A**, and the need for a large population of particles leads to long computation times.

### 3.6 A composite method for mapping

To exploit both the high time-efficiency of natural continuation and the global search capability of the particle swarm method, we develop a composite method that combines both approaches. An algorithm in pseudocode is shown in Appendix A.3. First (lines 1-12), we apply natural continuation starting from the simple-harmonic solution, as per §3.2. If, or when, the continuation step fails to find a solution, the algorithm switches to the particle swarm method for this step, using a population of randomly distributed initial guesses. If the particle swarm method still fails to find a solution, this frequency is considered the end of the contiguous resonant band and is labelled as the boundary frequency, Ω_*B*,up_ for the up and Ω_*B*,down_ for the down (lines 23-24). As with numerical continuation, separate upward and downward searches are performed, and their results are combined to obtain the complete resonant band. As an additional optional extension, to explore possible non-contiguous solution regions, we use the initial simple-harmonic solution as the sole initial guess to explore the remaining region of the search space (lines 25-30).

## 4 Results

### 4.1 Comparison of mapping methods

We test the three mapping methods developed over §§3.4-3.6 on the linear PEA flight motor, Eq. (3), with four waveform spectral terms (*N* = 4) and discretised dimensionless time of length 500 (*L* = 500), and considering also the size of the particle population (*M*) for PSO. Results are illustrated in Fig. 2, and quantified by the percentage area, *S*_*m*_, of search space, Ω ∈ [0, 2*ω*_*p*_], *ζ* ∈ [0, 1], over which a solution is to be found. The associated wall-clock computational time, *T*_*c*_ is also listed, based on computation on an x64-based Laptop equipped with Intel Core(TM) Ultra 5 235U (2000 Mhz) and 32 GB RAM. As can be seen, natural continuation identifies a continuous band with by far the shortest computational time, but the resulting band is incomplete due to premature failure of continuation steps. The particle swarm method explores the entire search space: results with two particles are superior to natural continuation, and the band appears complete with one hundred particles—though the computational time of the particle swarm method with one hundred particles is prohibitive. By contrast, the composite method achieves the same high-fidelity mapping as the prohibitive particle swarm method, but at only 3% time cost. This composite method is adopted for following computation. Note, however, one small but physically interesting feature of Fig. 2: the prohibitive particle swarm method identifies a single isolated solution at high damping (*ζ* = 1) and low frequency (Ω*/ω*_*p*_ ≈ 0.25), on the border of the search space. This is not numerical error, but an isolated band-type resonant state. In §4.2 we will explore this feature more fully.

**Figure 2:**
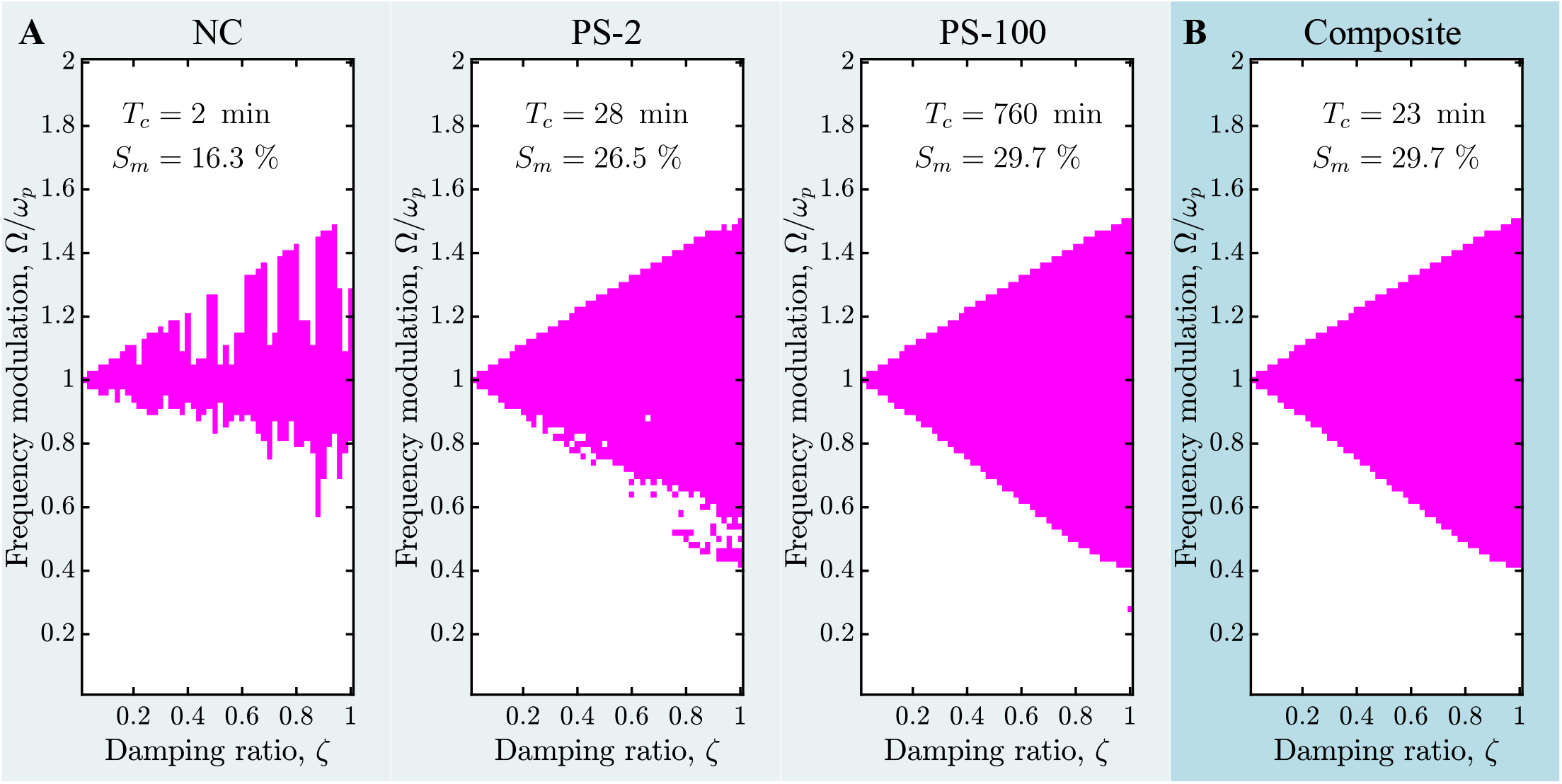
Performance of our suite of methods for mapping band-type resonance, applied to the linear PEA motor. **A.**Different single methods: NC, natural continuation; PS-2, particle swarm with 2 particles; and PS-100, particle swarm with 100 particles. **B**. A composite method, representing a cost-effective approach to mapping the complete contiguous resonance band.

With the composite method established as our mapping method of choice, we must then determine the appropriate number of waveform spectral terms (*N*)—in essence, a mesh convergence question. Mapping results by increasing number of spectral terms are presented in Fig. 3A for the linear PEA motor, with the solution area, *S*_*m*_, and computation time, *T*_*c*_ reported. We consider *N* = 6 to be a convergent spectral mesh: the boundary has reached a convergent area, and indeed the identified band under *N* = 6 is slightly wider than that under *N* = 7, because of the increasing likelihood of local optimisation non-converge in the larger parameter space. *N* = 6 is also significantly more time-efficient.

**Figure 3:**
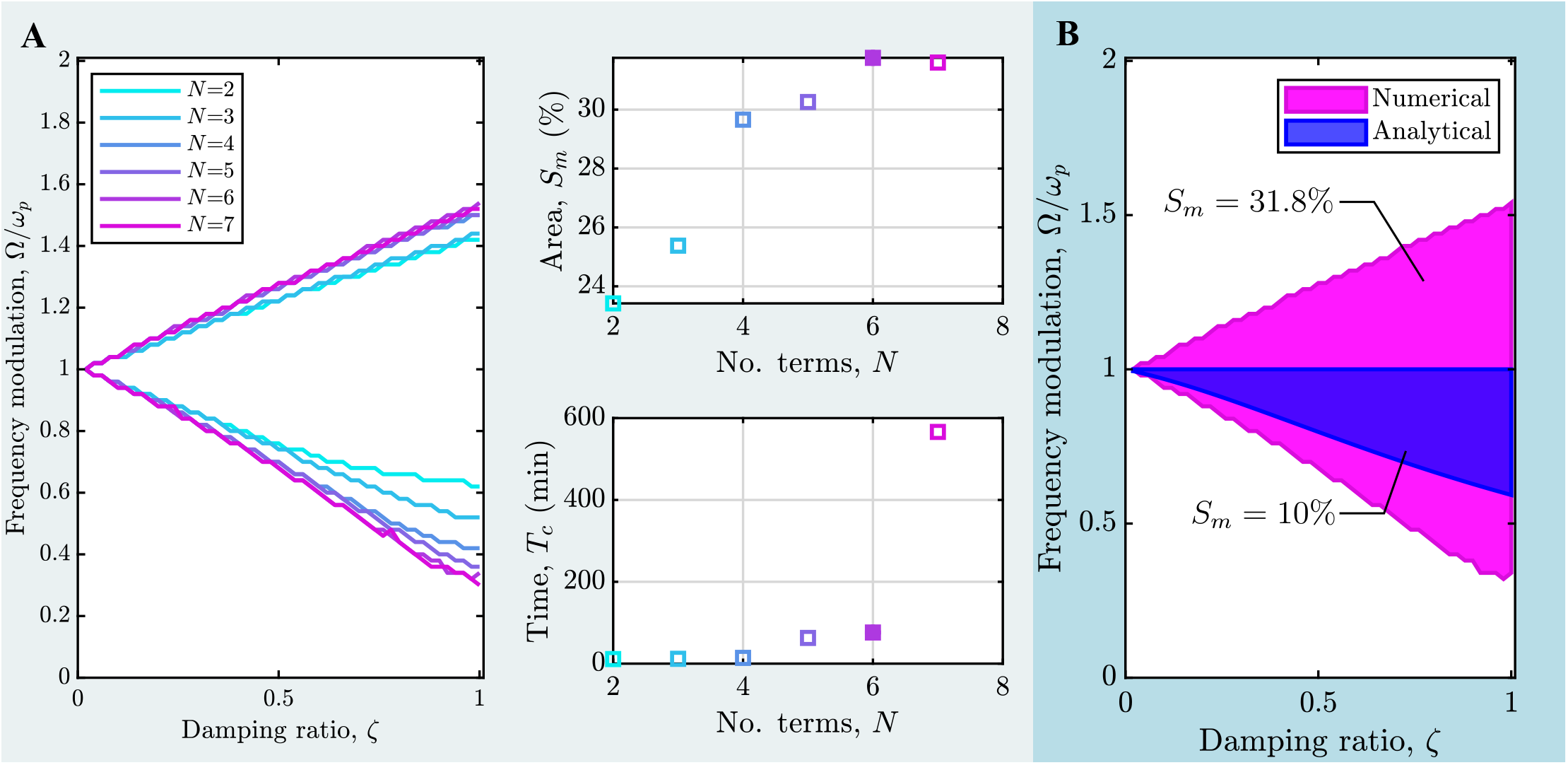
Mapping results for the linear PEA motor.**A.**The contiguous resonance band for increasing waveform generality, *viz*. number of spectral terms, *N*. Based on these results, we consider the method to be convergent to the complete results for smooth symmetric waveforms by *N* = 6. **B**. A comparison between our numerical mapping results and the previously analytical estimate of Eq. 7. Our results indicate that the resonance band is substantially larger than this previous estimate.

Fig. 3B compares these resonance band maps to the only previous results for this system, the analytical bound of Eq. (7) [24]. These numerical results advance on this previous result in two key ways: (**i**) they provide maps of the space of upward frequency modulation, Ω/*ω*_*p*_ > 1, and (**ii**) they provide maps which are not constrained by analytical approximation, but (we expect) approach the exact boundary of the resonance band for symmetric waveforms—a boundary which, as far as we know currently, is not computable analytically. The numerical maps expand on the analytical estimates by a factor of 3 in total resonant band area, or a factor of 1.5 considering only downward frequency modulation.

### 4.2 Band boundary features

These linear system results also reveal several interesting characteristics of the boundary of the contiguous resonance band—the extremal states, in fundamental frequency, that still maintain energy resonance. Fig. 4 presents the properties of the upper and lower boundaries of the resonance band, including the displacement *x*(*t*), force *F* (*t*), and power *P* (*t*) histories, normalised by their respective maxima for comparison. Across motor damping levels (*ζ*), these boundary states show the remarkably similar features. They are all characterised by intermittent power (and forcing) waveforms–that is, regions of high power and force output interspersed with regions of zero power and force output. This intermittency is tied to the direction of modulation: for upward modulation, this corresponds muscle forcing is concentrated in the first halves of the downstroke and upstroke; for downward modulation, it is concentrated in the second halves of the downstroke and upstroke. Tracking the points of peak muscle power output, *i*.*e*., requirement, more closely (Fig. 4B), we observe a curious property. For upward modulation, the timing of peak muscle power requirement is almost constant over damping ratio—indeed, the entire waveform remains almost identical. For downward modulation, however, the timing changes appreciably with damping ratio, nearing the stroke extrema as damping ratio increases.

**Figure 4:**
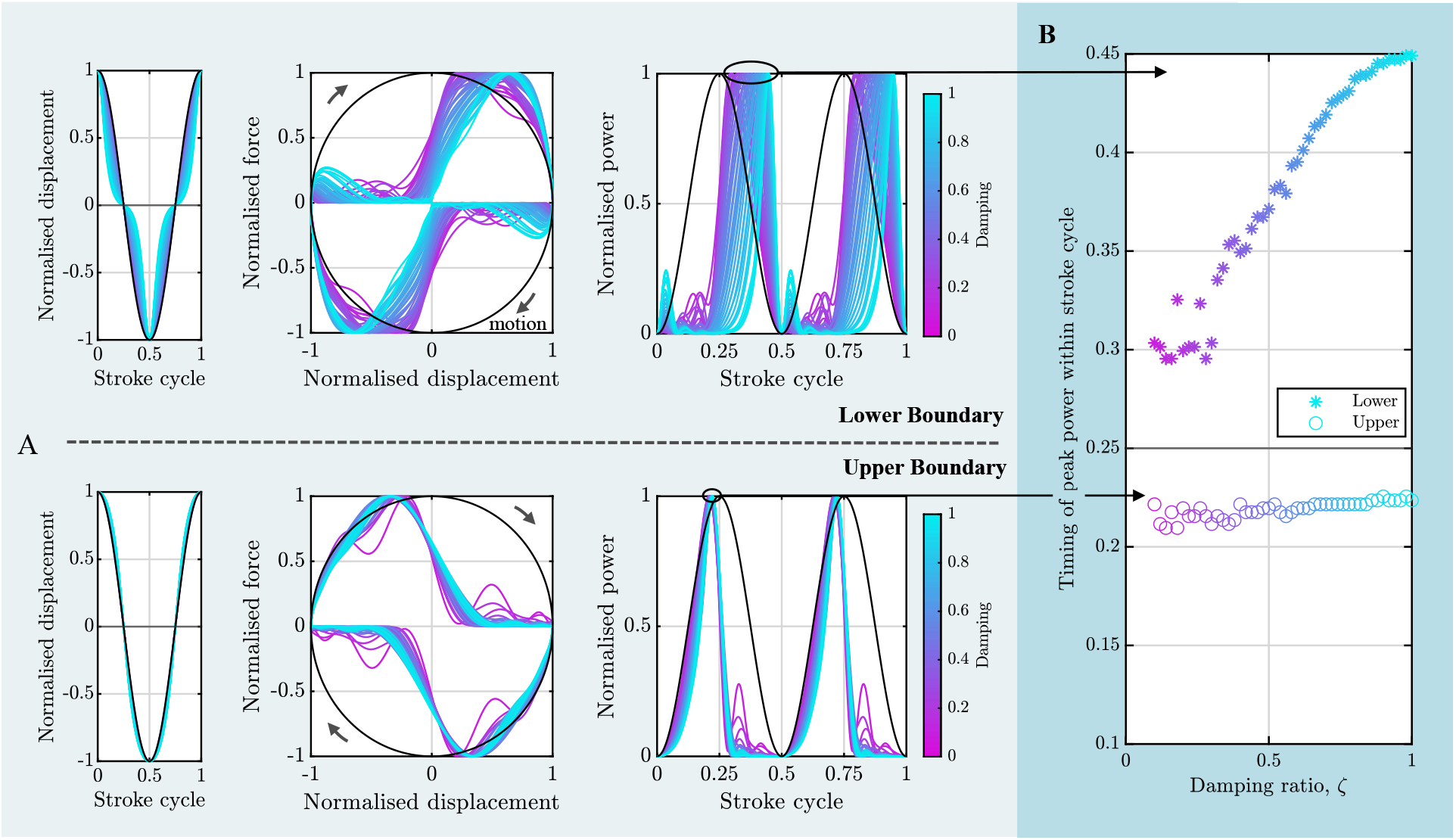
Characteristics of the boundary of the contiguous resonance band in the linear PEA motor.**A.**Normalised displacement, force, and power time histories for the upper and lower boundaries. Black lines represent simple-harmonic energy resonance, for comparison. **B**. Tracking the timing of the peak power within the stroke cycle, over the lower and upper boundaries. Curiously, the timing of the peak power on the lower boundary is nearly constant, whereas the timing on the upper boundary shows an increasing trend with damping ratio. Both trends suggest simple control strategies for energy-efficient wingbeat frequency modulation.

Both these trends provide a possible pathway to simple control strategies for FW-MAVs that preserve energy efficiency under wingbeat frequency modulation, and in the case of upward modulation from the natural frequency, it may not be necessary to know the level of damping in the system in order to achieve effective control. These trends also shed light on the action of the primary flight muscles in insect flight. Muscle force timing is a key mechanism for insect flight control [49], and our results indicate that this timing could itself act as a mechanism for energy-efficient wingbeat frequency modulation. This could occur outside the context of flight control: *e*.*g*., in the drosophilid sine song [50], a motor oscillation during courtship that occurs significantly below (≈ 70%) the wingbeat frequency; or in the context of results by Wold *et al*. [51] indicating that the wingbeat frequency of several insect species exceeds the thoracic natural frequency—both states might be controlled by muscle timing to remain within the resonance band.

Going further than these contiguous boundaries, we also explore the isolated solution at low frequency identified in Fig. 2. Fig. 5 presents an evaluation of the efficiency metric *η*, Eq. (6), over the search space, Ω ∈ [0, 2*ω*_*p*_], *ζ* ∈ [0, 1], under two conditions: (**i**) optimal waveforms from our method, including both band-type resonant states, *η* = 1, and others, *η <* 1; and (**ii**) simple-harmonic waves, representing frequency modulation with no waveform optimisation. Several features may be observed.

**Figure 5:**
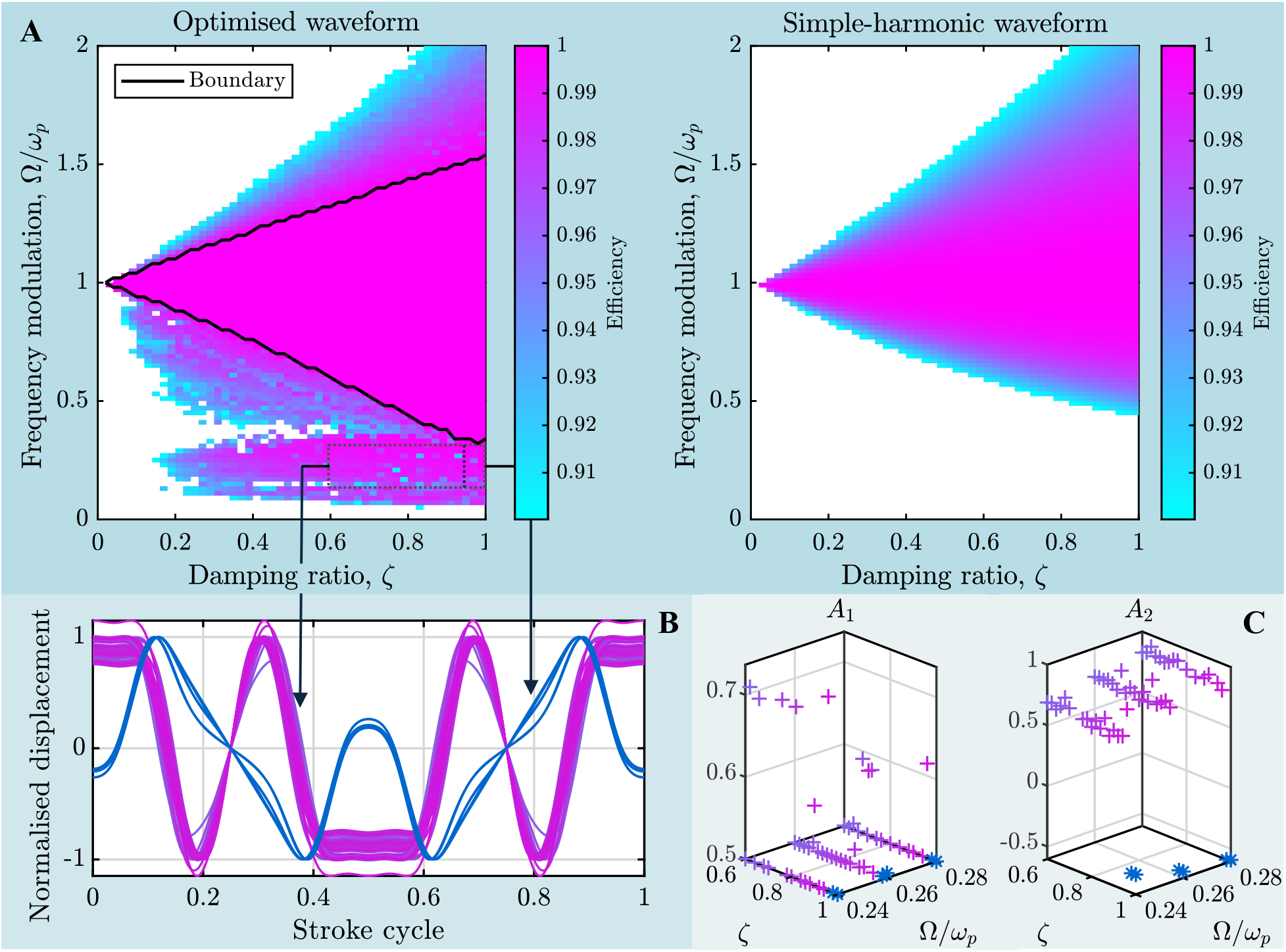
An isolated low-frequency high-efficiency region, described for the linear PEA motor.**A.**Efficiency comparison between under a simple harmonic waveform and a multi-harmonics sinusoidal waveform. An isolated high efficiency region emerges at high damping and low frequency under the multi-harmonic waveform. **B**. Overlaid displacement waveforms for the two marked subregions within this high-efficiency region, coloured according to subregion. Each subregion has a consistent and distinctive waveform, which is strongly multiharmonic. **C**. The first two spectral coefficients (*A*_1_, *A*_2_) for these distinctive waveforms, coloured according to region. Most, but not all, of these waves lie on the constraint boundary for aliasing, | *A*_1_ | *>* 0.5. This isolated high-efficiency region relies on higher harmonics high efficiency, but it still contains a significant spectral component at the fundamental frequency. Interestingly, the spectral coefficient *A*_2_, which is significant or dominant to *A*_1_, does *not* alias onto the natural frequency, *ω*_*p*_, but rather onto ≈ 0.75*ω*_*p*_.

1. An isolated high-efficiency region—distinct from the main band around the natural frequency—appears under waveform optimisation. This region appears around the aliasing region, *cf*. §3.4, and we surmise that it arises because of the coincidence of higher harmonics with the natural frequency. However, these near-solutions are not complete aliases, as illustrated in the waveform and spectral coefficient values, Fig. 5B-C. They are rather strongly multiharmonic waves, with significant spectral amplitudes both at the fundamental and the next harmonic.
2. As identified previously by Pons & Beatus [9], energy-efficiencies under even simple-harmonic wingbeat frequency modulation remain high—this is because the energy transfer function is significantly more lenient (in this sense) than the classical transfer functions. Waveform optimisation converts these near-resonant states to true energy-resonant states, and also makes significant improvements to downward frequency modulation. An implication here is that band-type resonance might be more effective for downward modulation— however, this might be a feature of the specific waveforms that we consider here (symmetric with only higher harmonics). As per (**1**), low-frequency solutions are assisted by aliasing of higher harmonics of the fundamental frequency, and so it appears likely that high-frequency solutions would be assisted by subharmonics of the fundamental. This is an interesting case for future study, which our numerical methods are able to tackle.

### 4.3 Influence of damping on band

Moving from the linear to the quadratic PEA motor, Eq. (4), we map the contiguous resonance band and compare the two motors in Fig. 6. We compare the two damping parameters, *ζ* and *B*, on a consistent basis via equivalent net power output at the simple-harmonic energy resonant state at *ω*_*p*_. This net power is a proxy for the level of damping in the system. As per Fig. 6, the width in frequency of the resonant band in the quadratically-damped motor is smaller than that of the equivalent linearly-damped motor. However, because of the stronger dissipation scaling in the quadratically-damped system, the available modulation in power throughput is greater. Given that this power throughput—rather than frequency, *per se*—determines the propulsive force generated by the motor, the increased modulation is likely a beneficial characteristics. This suggests that insects benefit from their approximately quadratic aerodynamic drag scaling, in terms of energy-efficient manoeuvring.

**Figure 6:**
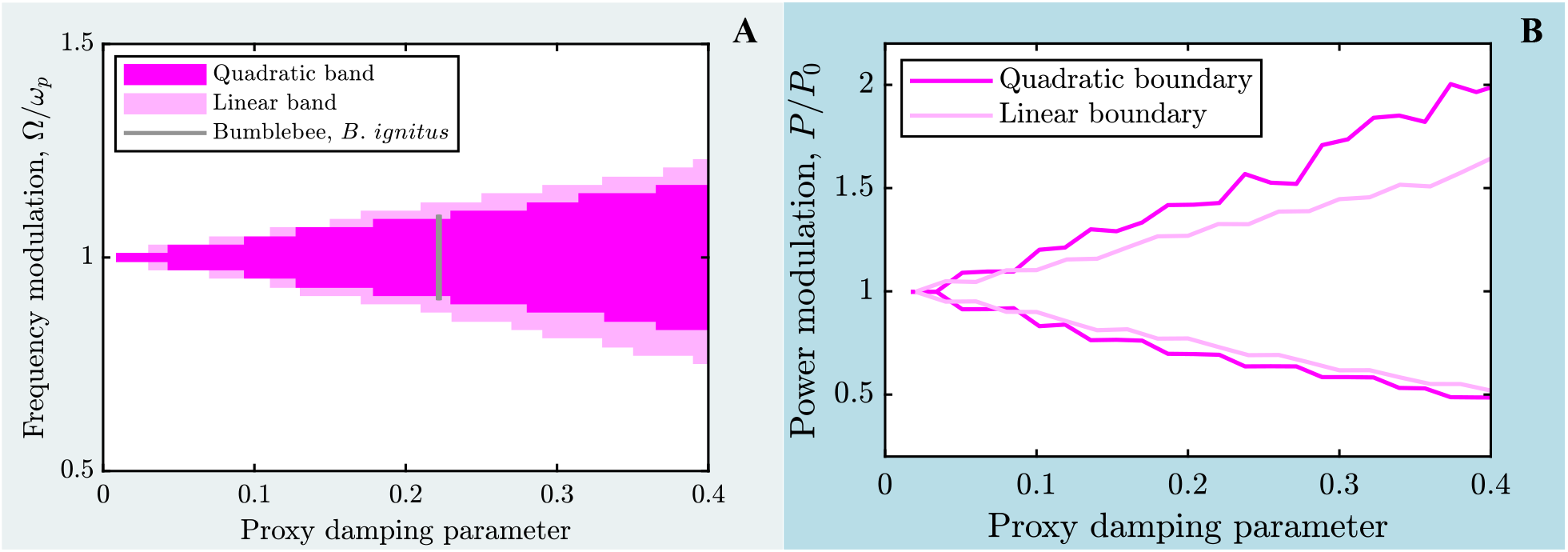
Resonance band by PEA motor dissipation scaling: linearly-vs. quadratically-damped. Maps of the resonance band are compared based on the equivalent net power output at the simple-harmonic energy resonant state at *ω*_*p*_ (the proxy damping parameter). While the width, in frequency (**A**), of the resonance band in the quadratic system is smaller than that in the linear system, the actual power modulation (**B**) available to the quadratic system is greater, particularly in terms of increased power output—a potentially advantageous feature of the stronger damping scaling.

Fig. 6 also includes a comparison between these resonance bands and reported frequency modulation ranges measured in the bumblebee *Bombus ignitus* [52]. Motor parameter values for this specie are identified in the Appendix A.4, and data by Iwamoto & Yagi [28] indicates PEA model applicability. Reported wingbeat frequency modulations are within the resonant band, indicating that these modulations can, in theory, be carried out without loss of efficiency via band-type resonance. This does not discount the possibility of other mechanisms at work during these modulations, *e*.*g*., effective natural frequency changes due to changes in muscle activation, but it does indicate that there is not necessarily any penalty, in mechanical efficiency, to performing these modulations without these other mechanisms.

### 4.4 Distributed elasticity: a moderate-risk, high-reward strategy

Having established the properties of the resonance band for the case of parallel-only elasticity, we can compare how distributed elasticity—between thoracic (parallel) and wing root (series) affects this band. We apply our composite mapping method to the HEA motor, Eq. (5). Fig. 7 illustrates a selection of results from this comparison.

1. Considering first the motor’s simple-harmonic behaviour, Fig. 7A.1 presents the efficiency (*η*) curves for simple-harmonic motion over wingbeat frequency (Ω) and motor damping (*B*), for an illustrative case of *α*_*r*_ = 1.5. In general, the motor has two simple-harmonic energy resonant states. At increasing damping, *B*, these two states converge: a lower (parallel-type) state persists at a constant frequency, and a higher (series-type) state drops towards the former. Notably, beyond a certain critical level of damping (dependent on *α*_*r*_), the lower energy resonance becomes extinct: at peak, *η <* 1, *i*.*e*. pseudo-energy resonance [9]. While the loss in efficiency is small, from the perspective of an insect or FW-MAV relying on self-oscillation—via asynchronous muscles or actuators [43, 53]—this is concerning. It is not a matter of simply supplying more energy to overcome the loss of efficiency: linear (or, linearised) self-oscillation requires an energy resonant state in order to occur [43], and the lack of such a state implies that the motor will not self-oscillate irrespective of the power input.
2. Considering next the motor’s band-type resonant states, Fig. 7A.2 presents our complete estimate of the resonance band for *α*_*r*_ = 1.5, accounting for the branches around both the lower and upper energy resonances via respective applications of our composite mapping method about these frequencies. The branch mapped solely via the lower energy resonance is indicated—this lower branch is then mapped across a range of *α*-values in Fig. 7B.1. We do not consider the higher branch to be biologically relevant: operating within it would imply large (supercritical) phase lags between muscle and wingbeat, which have not been observed in any species of insect. For any particular elasticity distribution (*α*_*r*_), the resonance band shows a three-step trend of equivalence-expansion-extinction. Initially, at low damping, the resonance band is equivalent in size to the PEA system: in this regime, the upper and lower branches do not interact. Then, beyond a certain critical level of damping (dependent on *α*_*r*_), the resonance suddenly and massively expands due to the branch interaction. Finally, beyond a second critical level of damping, the resonance band is suddenly extinguished: physically, it becomes impossible to transmit enough sufficient energy through the wing root elasticity to overcome the strong wing damping. For a very stiff wing root (*e*.*g*., *α*_*r*_ = 3), this final extinction lies well beyond the damping level of any insect; but for a softer wing root (*e*.*g*., *α*_*r*_ = 1), many species would be subject to resonance extinction.
3. The same three steps of equivalence-expansion-extinction can be seen in the behaviour of the band for any particular damping under changing *α*_*r*_, *i*.*e*., the tuning of wing root flexion for a particular insect species. Initially, for a very stiff wing root, the motor shows PEA-equivalent behaviour; then, below a critical stiffness, the resonance band suddenly expands; and finally, below a second threshold, the resonance band is extinguished. Considering this from the perspective of an evolutionary optimisation of an insect species’ wing root, the window of beneficial stiffnesses is relatively large, and the primary concern would be to avoid straying into the extinction region—only a moderate risk. But the (contextual) reward—in terms of the expansion of the window of efficient wingbeat frequency modulation, and/or the ability to operate at wingbeat frequencies higher than the thoracic natural frequency—is potentially large. Fig. 7B.2 compares data on observed levels of wingbeat frequency modulation [52, 54, 55], and estimated elasticity distributions, for three insect species: the bumblebee *Bombus ignitus*, the fruitfly *Drosophila melanogaster* and the hawk-moth *Agrius convovuli*. The estimation process for species damping ratio is detailed in Appendix A.4, and for elasticity distribution, in Appendix A.5. For all three species, observed wingbeat frequency modulation lies within the resonance band—indicating that, in theory, this modulation could be performed without loss of mechanical efficiency. Both *Drosophila melanogaster* and the hawkmoth *Agrius convovuli* appear to reap the awards of distributed elasticity (*α*_*r*_ ≈ 1.5), via significant expansion of the resonance band (by a factor of ≈ 2). Both species could afford to have wing roots approximately 30% softer, *e*.*g*., via wear or developmental variation, before resonance extinction: illustrating that distributing elasticity over the motor can bring rewards at only moderate risk.

**Figure 7:**
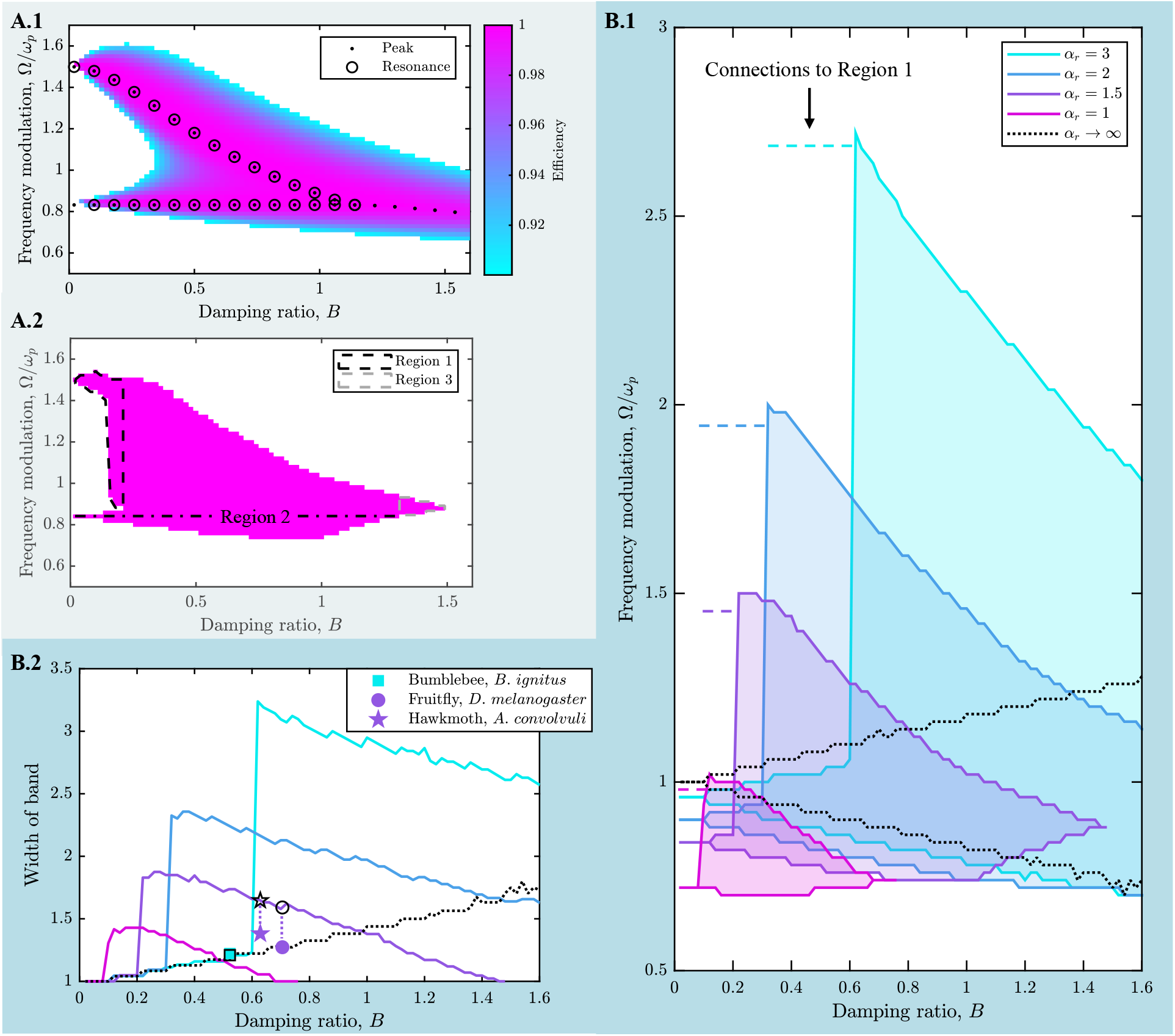
Influence of wing root flexion on the resonance band: mapping results for the HEA motor. **A.1** Efficiency map, for *α*_*r*_ = 1.5 under a simple harmonic waveform, with efficiency (*η*) peaks illustrated as a function of *B*. Where these peaks reach *η* = 1, they are energy-resonant. The extinction of energy resonance at sufficiently high damping can be observed. **A.2** Map of band-type resonance for *α*_*r*_ = 1.5. To obtain this map we apply our suite of methods over three regions: the composite method around the upper (Region 1) and lower (Region 2) energy resonances; and the particle swarm method around the resonance extinction point (Region 3). **B.1** Maps of band-type resonance over *B* and *α*_*r*_, for Regions 2-3. The three-step process of equivalence (to PEA), sudden and massive expansion, and then eventual extinction can be seen. **B.2** The width of the resonance mapping, *i*.*e*., the ratio between the maximum and minimum Ω at which band-type resonance exists, representing the maximum available wingbeat frequency modulation factor under band-type resonance. Wingbeat frequency modulation factors observed in three insect species (coloured symbols) are compared to predictions of the maximum available factor (black symbols).

## 5 Discussion and conclusion

### 5.1 Methodological advances

In this work, we develop the first methods capable of mapping the complete space of band-type resonant states in general dynamical systems. The application of these methods to models of the insect flight motor yields insight into insect functional morphology and FW-MAV design—as we discuss further in §§5.2-5.3. Their relevance as a methodological advance is twofold.

1. They allow mapping of the complete set of band-type resonant states, rather than only an analytically convenient subset as per [24, 27]. This allows us to make fair and quantitative comparisons between systems and thereby to assess whether system parameters—such as the distribution of elasticity within an insect flight motor—alter the system’s resonant band beneficially or detrimentally. It also allows us to identify deep properties of this band, such as the power timing characteristics of the boundary, Fig. 4.
2. These methods are applicable to complex systems that are beyond analytical treatment. This applicability is a key step to allow future identification of whether or not insects actually use band-type resonance—for instance, by comparing insect kinematics under flight control to the resonance band for high-fidelity computational models of fluid-structure interaction in the flight motor [56]. Such an approach, while computationally expensive, is now enabled.

### 5.2 Implications for insect functional morphology

Insects show great diversity in the spatial distribution of flight motor elasticity: over muscle, tergum, pleural wall, wing root, and wing. The adaptive value—typically pertaining to general flight efficiency or performance—of many of these individual elasticities has been identified or hypothesised [6, 12, 32, 57, 58]. This adaptivity raises the broader question of why there are differences in distribution—*e*.*g*., if wing flexion improves flight efficiency [57], why is it only observed in some insect species? Furthermore, there is also a certain redundancy in the adaptive value of these different elasticities—*e*.*g*., a resonant wingbeat could be achieved via elasticity pleural wall elasticity or wing root elasticity; both are not required [9].

Here, we identify previously-unrecognised adaptive value to distributing the elasticity across the flight motor: this distribution, if tuned to the level of damping in the motor, can massively increase the window of wingbeat frequencies over which the motor is efficient. However, if this distribution is mistuned, then this window of high efficiency can vanish. These pair of mechanical phenomena offer conjectural explanations for the diversity in insect motor distribution. We conjecture that distributed elasticity is particularly important to insects that require significant flight manoeuvrability, and, in particularly, the ability to rapidly modulate overall lift force for sudden ascending or descending flight. The well-studied fruit fly *D. melanogaster* provides a single case to support this conjecture: similar to other dipterans, *D. melanogaster* is adapted to manoeuvrable flight via the dipteran halteres, and to rapid takeoff [59]. *D. melanogaster* is one species for which we have clear evidence of significant wing root elasticity, and, as per our analysis, the presence of this elasticity assists it in efficient wingbeat frequency modulation.

We conjecture also that distributed elasticity may also be valuable to insects that—for as-yet unknown reasons—may operate at wingbeat frequencies significantly exceeding the thoracic natural frequency, *cf*. Wold *et al*. [51]. Our results reveal the counterintuitive principle that higher wingbeat frequencies are enabled by softening, rather than stiffening, a structure: the base of the wing. The mechanism behind this principle is that, when the natural frequencies of the wing and thorax start to approach each other—within a factor of roughly 2*×*—the space of band-type resonance extends between the two resonances, enabling self-oscillation and/or an energy-efficient wingbeat well above the thoracic natural frequency. Future comparison of insect wingbeat kinematics with resonance band predictions for high-fidelity computational fluid-structure interaction models, as we discuss in §5.1 could help test this pair of conjectures.

### 5.3 Implications for the design of FW-MAVs

Designing distributed elasticities into FW-MAV structures offers the possibility of efficient wingbeat frequency modulation—a feature particularly relevant to FW-MAVs with wingbeat amplitudes that are fixed by kinematic mechanisms [60]. The power timing characteristics identified in Fig. 4B indicate that it may be possible to design simple control strategies that take advantage of this efficiency. Designing distributed elasticity into an FW-MAV could potentially be achieved with minimal structural changes, by simply by softening the wing root enable flexion. While accessible, there are also risks associated with this approach, if the wing root flexion is mistuned. Softer wing root structures have detrimental effects on flight efficiency—and could potentially confound novel FW-MAV designs relying on self-oscillation [53].

## Supporting information

Code for the work

## Ethics

This work did not require ethical approval from a human subject or animal welfare committee.

## Data accessibility

Data and code used in this study are attached as supplementary material.

## Declaration of AI use

We have not used AI-assisted technologies in creating this article.

## A.1 Mapping algorithm based on natural continuation

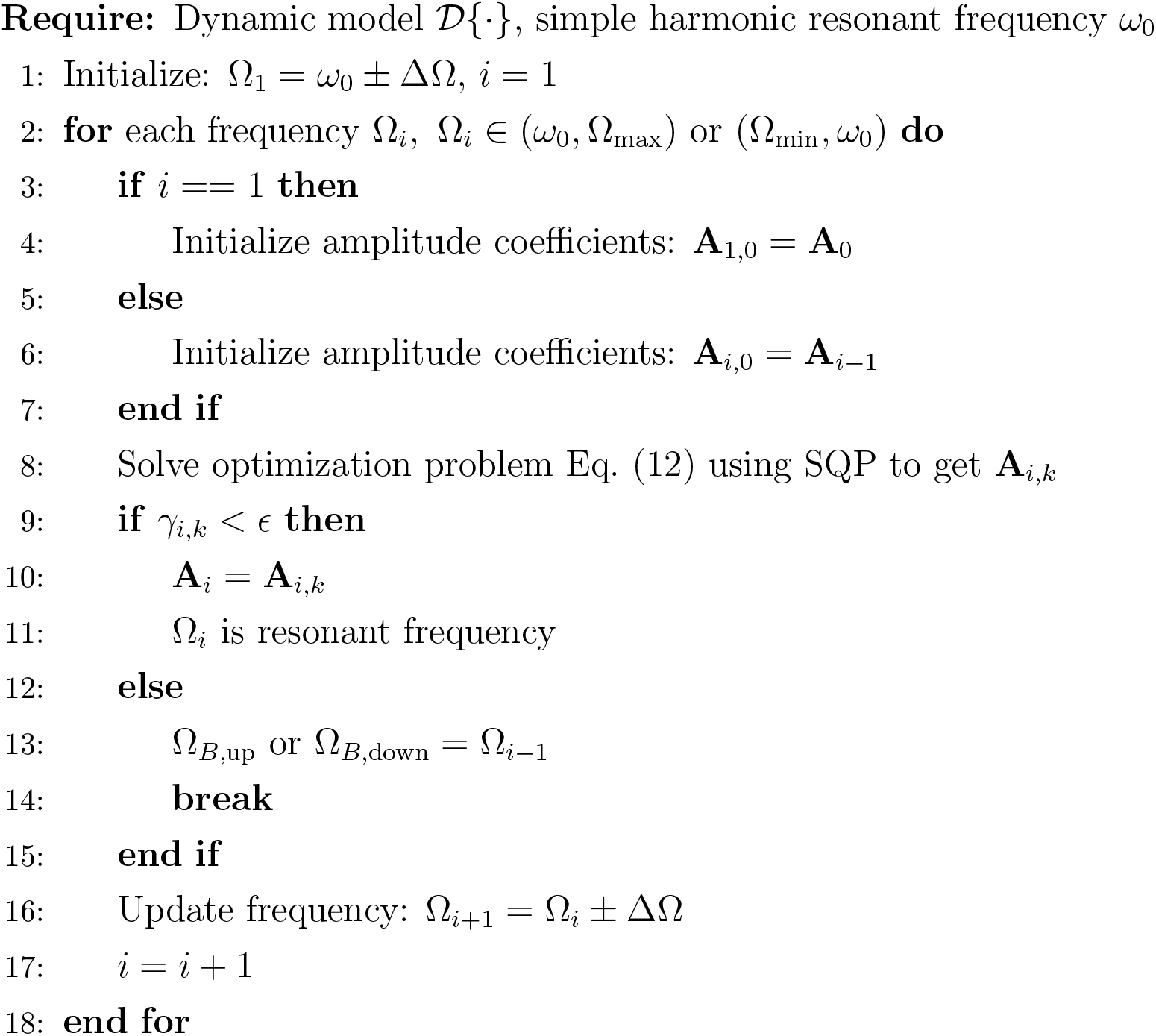

*Note: ω*_0_ is a resonant frequency with a simple harmonic state. For the linearly or quadratically damped PEA motor, *ω*_0_ = *ω*_*p*_, while in the HEA motor, *ω*_0_ can be found (if it exists) via a numerical parameter sweep over the frequency and the solution may not be single. The flow chart applies to both upward search—(*ω*_0_, Ω_max_)—by increasing the step *ω*_0_ + ΔΩ and Ω_*i*_ + ΔΩ, and downward search—(Ω_min_, *ω*_0_)—by decreasing the step *ω*_0_ − ΔΩ and Ω_*i*_ − ΔΩ.

## A.2 Mapping algorithm based on a particle swarm method

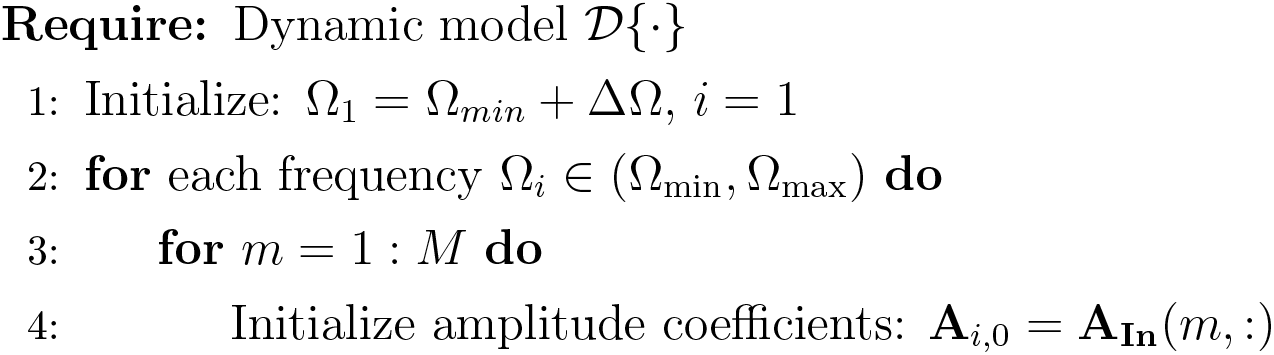
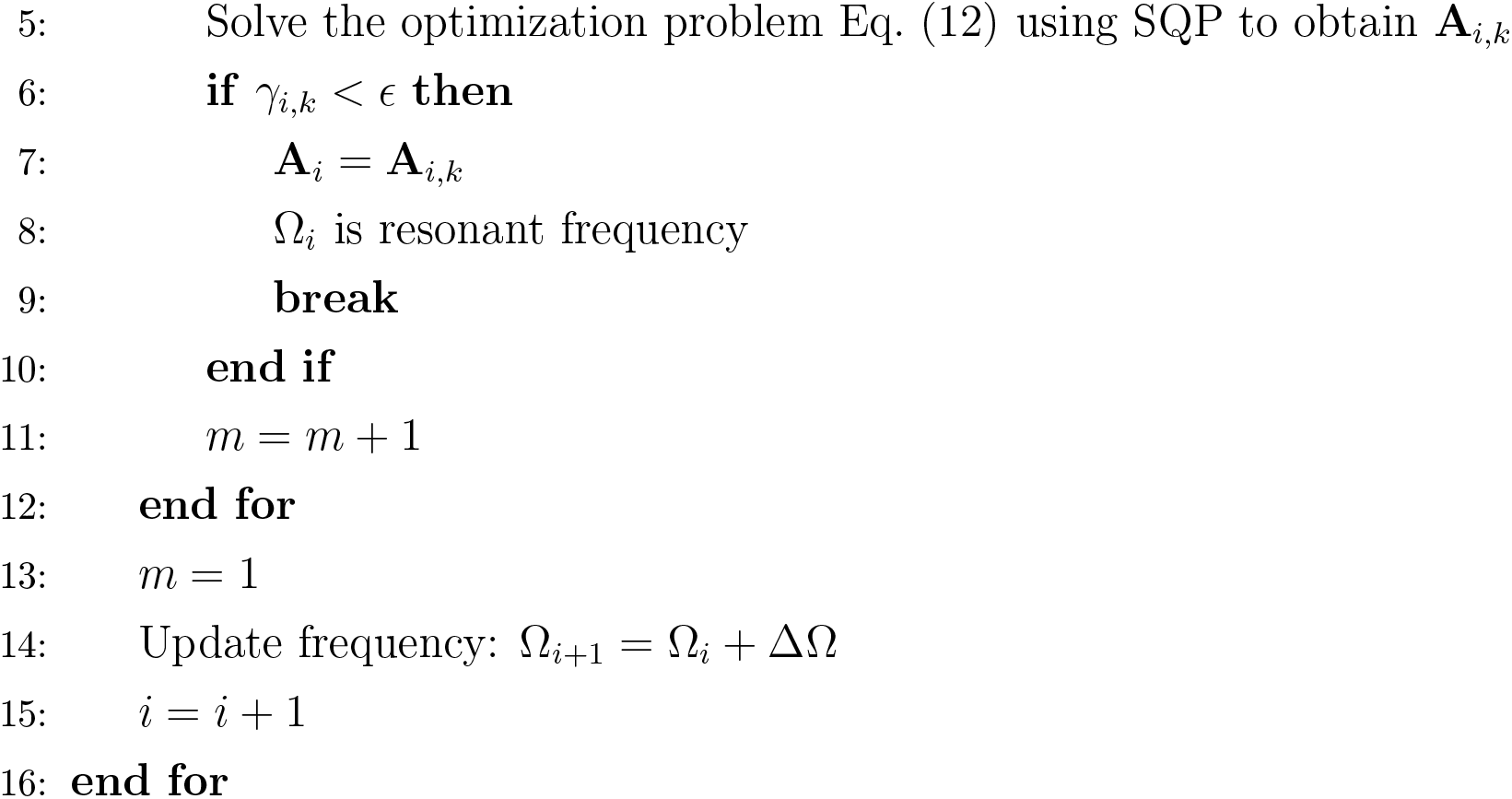

## A.3 Composite mapping algorithm

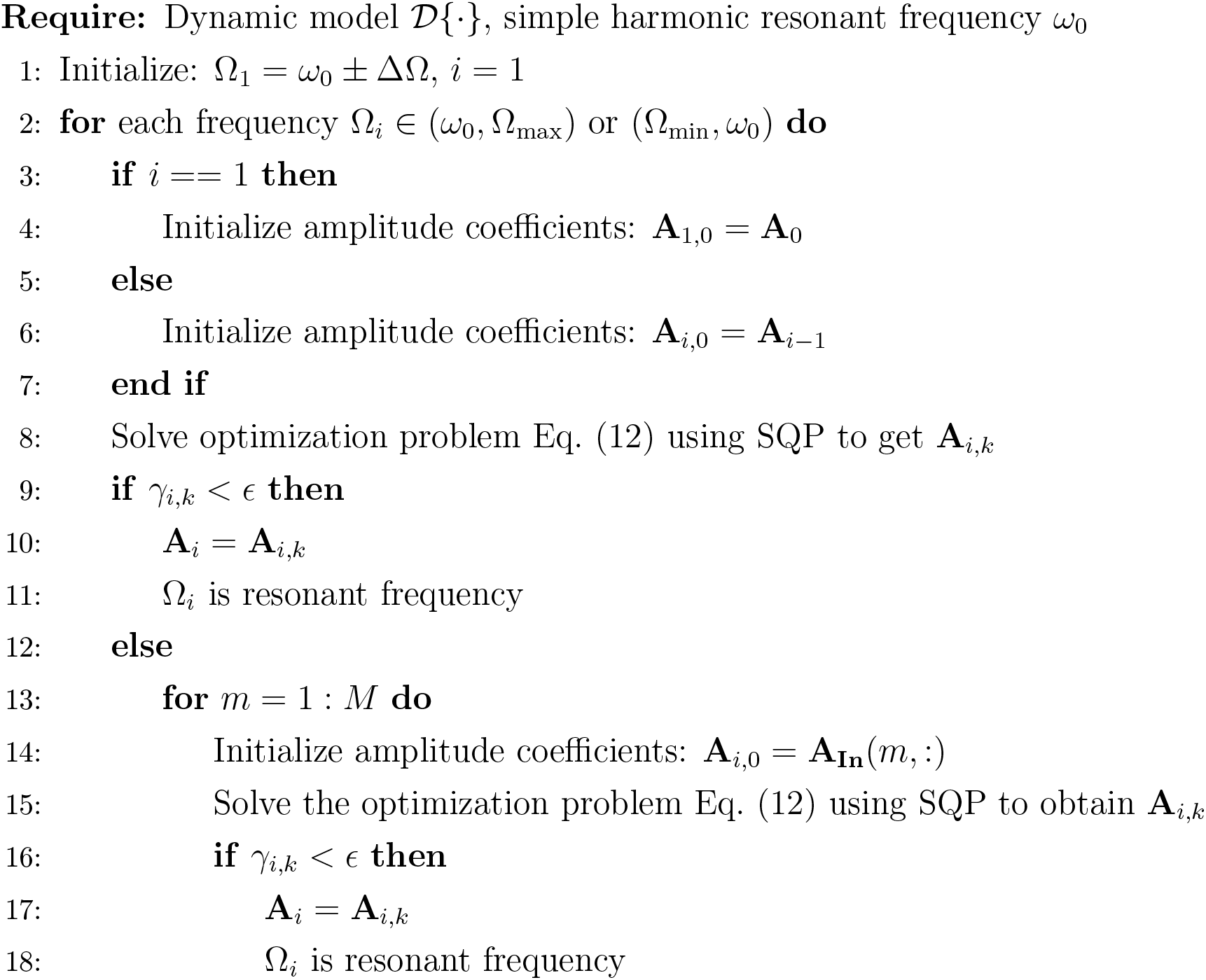
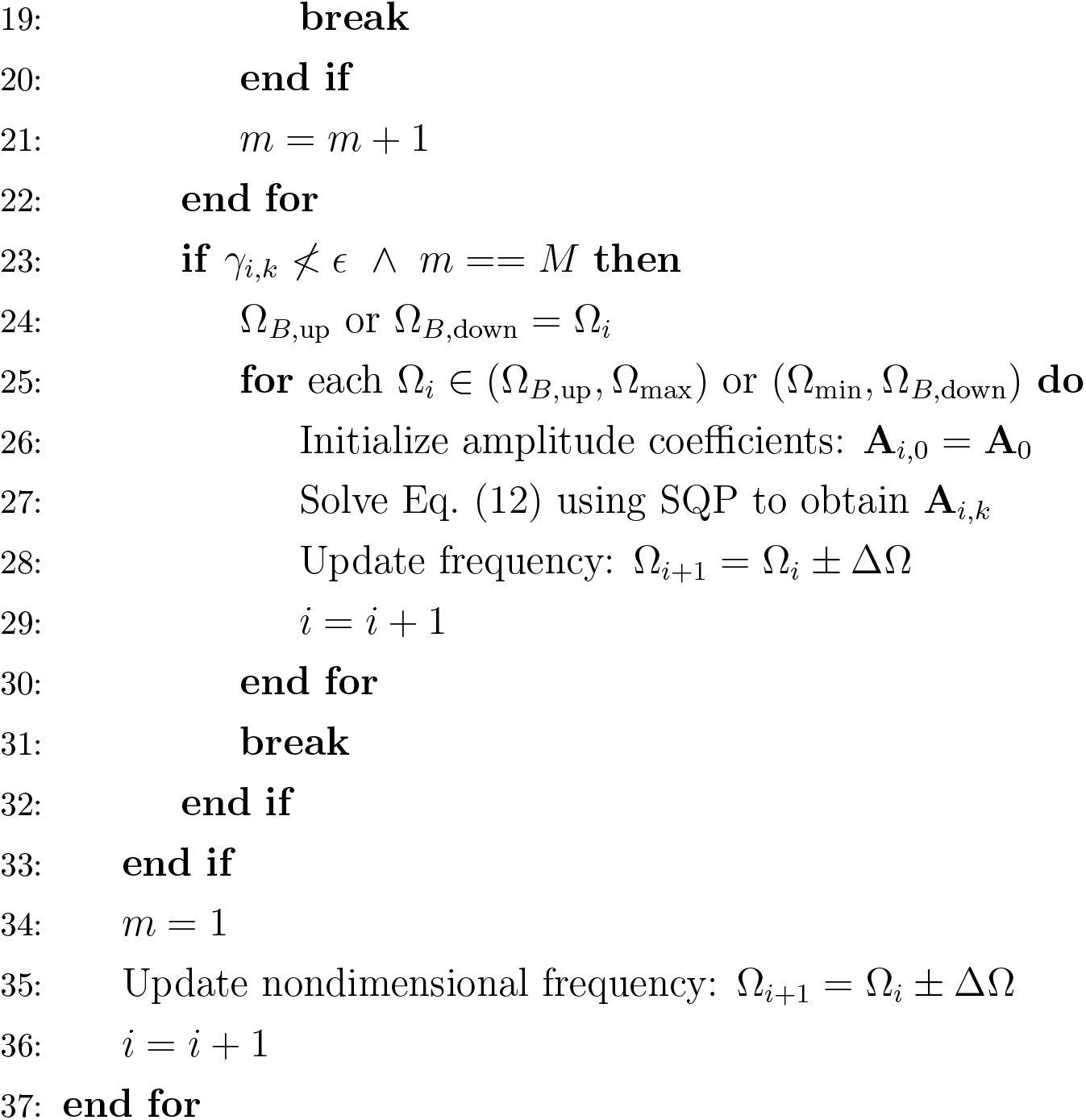

*Note: ω*_0_ is a resonant frequency with a simple harmonic state. For the linearly or quadratically damped PEA motor, *ω*_0_ = *ω*_*p*_, while in the HEA motor, *ω*_0_ can be found (if it exists) via a numerical parameter sweep over the frequency and the solution may not be single. The flow chart applies to both upward search—(*ω*_0_, Ω_max_), (Ω_*B*,up_, Ω_max_)—by increasing the step *ω*_0_ + ΔΩ and Ω_*i*_ + ΔΩ, and downward search—(Ω_min_, *ω*_0_), (Ω_min_, Ω_*B*,down_)—by decreasing the step *ω*_0_ − ΔΩ and Ω_*i*_ − ΔΩ.

## A.4 Parameter estimation: quadratic damping ratio, *B*

A dimensionalised version of the quadratically-damped flight motor, Eq. (4), may be expressed:

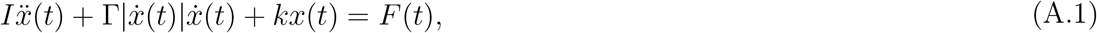

where *x*(*t*) is the wing stroke angle, *I* is the wing inertia, Γ is the coefficient of aerodynamic damping due to drag, and *k* is the linear or linearised thoracic stiffness. Comparing Eq. (A.1) with its dimensionalised form, Eq. (4), we see that *B* = Γ*/I*. This model has been considered previously by Wold *et al*. [61] and Lynch *et al*. [46], among others, who define the Weis-Fogh number, *N*_WF_, as the ratio of peak inertial to peak aerodynamic force over a wing stroke for sinusoidal motion:

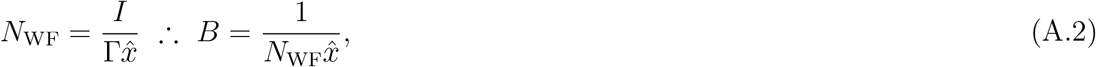

where 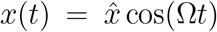, and for our resonance band analysis, we have taken 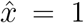 for normative hovering flight.

Wold *et al*. [61] identify *N*_WF_ in terms of the classical wingbeat-averaged inertial 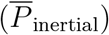 and aerodynamic 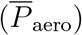 power measures [62], as:

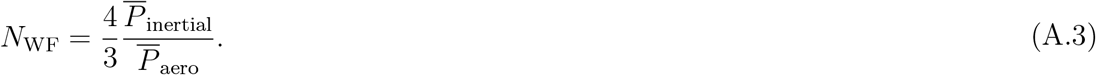

Results for these power measures allow computation of *N*_WF_ and thereby *B*. Table A.1 illustrates our estimations for three species: the bumblebee *Bombus* spp., fruit fly *Drosophila melanogaster*, and hawkmoth *Manduca sexta*. Specially, we generalize the damping data of bumblebee *Bombus* spp. to bumblebee *Bombus ignitus*, and that of hawkmoth *Manduca sexta* to hawkmoth *Agrius convovuli*, due to the data availability and considering their biological similarities.

**Table A.1:**
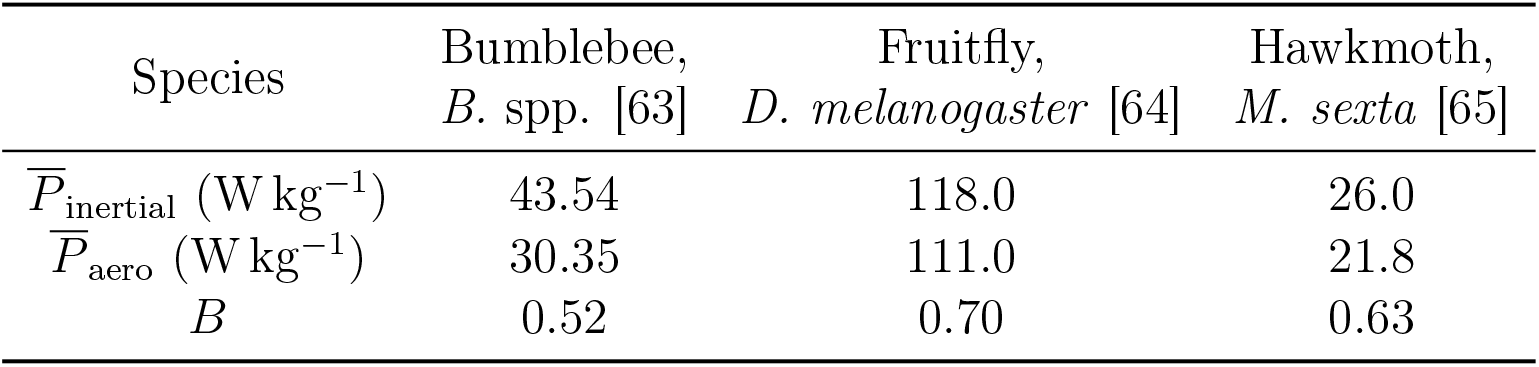
Estimated flight motor quadratic damping ratio for three insect species.

## A.5 Parameter estimation: elasticity distribution *α*_*r*_

We identify the elasticity distribution parameter, *α*_*r*_, indirectly based on observations of phase difference in motion across the flight motor. Based on the HEA dynamics of Eq. (5), the relationship between the muscle displacement *u*(*t*) and the wing displacement *x*(*t*) is:

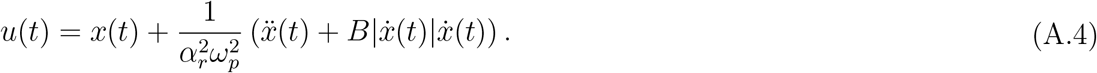

Assuming that the motor is operating with simple-harmonic output at the parallel natural frequency, *x*(*t*) = sin(*ω*_*p*_*t*), we may compute the associated input, *u*(*t*), which is multiharmonics due to the nonlinear damping. We apply the fast Fourier transform (FFT) to this wave to obtain a decomposition 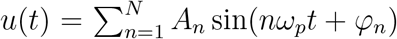. To measure the phase lag between *u* and *x*, we retain only the fundamental component *û*(*t*) = *A*_1_ sin(*ω*_*p*_*t* + *φ*_1_) and extract *φ*_1_.

We may then compare this phase, *φ*_1_, which depends on *B* and *α*_*r*_, to measurements of the wing-thorax phase difference across the literature. We take the meta-analysed data of Pons & Beatus [9], deriving from X-ray diffraction [28, 66] and laser profilometry [32] studies; as well as the estimates of *B* made in Appendix A.4. Note that the phase difference of bumblebee *Bombus* spp. from [9] is generalized to bumblebee *Bombus ignitus* due to the data availability and considering their biological similarities. Via numerical optimisation, we identify *α*_*r*_ such that the model phase estimate matches these experimental phase measurements: results are illustrated in Fig. A.1. We note that this estimation relies on two assumptions. (**i**) We have assumed that the wingbeat frequency coincides with the parallel natural frequency, *ω*_*p*_, which may not be the case—*e*.*g*., if an insect instead operates much higher its resonance band, cf. Fig. 7 and Wold *et al*. [51]. If the relationship between the wingbeat frequency and *ω*_*p*_ is known, the identification can be adapted. (**ii**) We have attributed all of the experimentally-measured phase difference to chordwise flexion at the wing root, *i*.*e*., *α*_*r*_. In the case of *D. melanogaster*, Melis *et al*. [10] provide direct observations of this flexion; but in *A. convovuli* it is less clear whether observed phase differences are due to wing flexion; a thoracic local mode; or both [9, 32]—further studies are required. In the case of *B. ignitus*, even though the observed phase lag is very small (≈ 3°), generally indicating PEA model applicability, the elasticity in the wing root is not completely negligible (*α*_*r*_ ≈ 3), and there is a significant expansion of the resonance band, as per Fig. 7. This may indicate that even these small phase lags may have significant function.

**Figure A.1:**
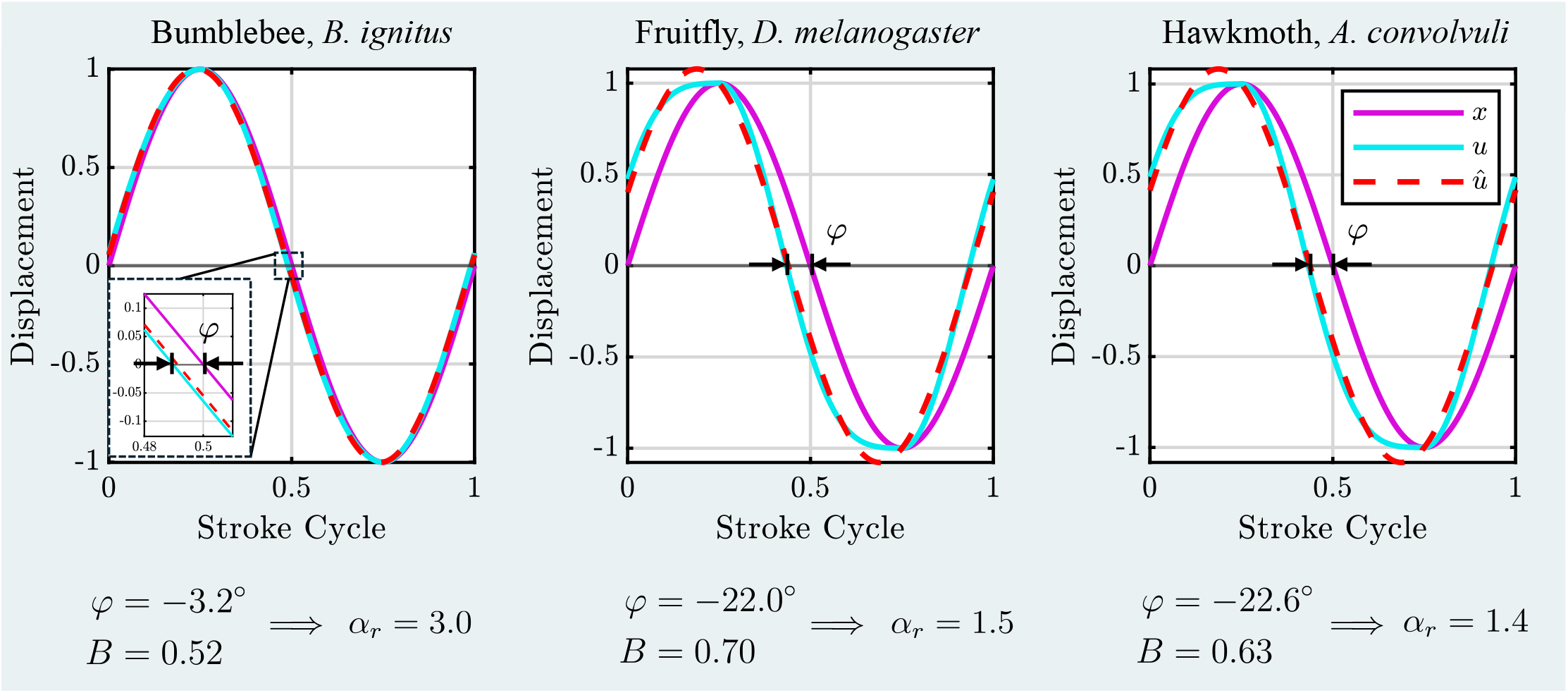
Estimating the motor elasticity distribution (*α*_*r*_) for three species based on reported wing-thorax phase data. *x* is the prescribed sinusoidal wing displacement; *u* is the corresponding muscle displacement, based on Eq. (A.4); and *û* is the fundamental frequency component of *u*.

