## Supplementary material for "Distributed elasticity: a high-reward, moderate-risk strategy for efficient control modulation in insect flight": Code for the work: composite.m

```
%%%%%%%%%%%%%%%%%%%%%%%%%%%%%%%%%%%%%%%%%
% Composite method for Mapping the space of band-type resonance in insect and insect-inspired
% Both the band and the non band will be explored
% Apply for the PEA system: Linear and quadratic damping
% Author: Liang Wang, Congxiao Zhang, Najmeh Asadimoghaddam, and Arion Pons
% Date: 18th February 2026
% Matlab version: R2023b
% -------------------------------------------------------------------------
clear all
close all
clc

% Numerical methods for computing band-type resonant states
% f = DDX + 2 * zeta * omega_0 * DX + omega_0^2 * X;
% p = f .* DX;
% We aim to find states with p>0
%% Graphics
% Colours, neon colourset:
NE_pi = [217 12  220]/255;
NE_cy = [4   236 239]/255;
NE_na = [75  88  167]/255;
NE_bk = [0   0   0  ]/255;
NE_wh = [1   1   1  ];
NE_gr = 0.3*[1 1 1];
NE_gm = 0.5*[1 1 1];
NE_gl = 0.7*[1 1 1];

NE_pibk = 0.7*NE_pi + 0.3*NE_bk;
NE_cybk = 0.7*NE_cy + 0.3*NE_bk;
NE_piwh = 0.4*NE_pi + 0.6*NE_wh;
NE_cywh = 0.4*NE_cy + 0.6*NE_wh;
NE_nawh = 0.6*NE_na + 0.4*NE_wh;

% Defaults:
set(groot, 'defaultAxesTickLabelInterpreter','latex');
set(groot, 'defaultLegendInterpreter','latex');
set(groot, 'defaultTextInterpreter','latex');
set(groot, 'defaultTextFontSize', 11);
set(groot, 'defaultAxesXColor', [0 0 0]);
set(groot, 'defaultAxesYColor', [0 0 0]);
set(groot, 'defaultAxesGridAlpha', 1);
set(groot, 'defaultAxesMinorGridAlpha', 1);
set(groot, 'defaultAxesGridColor', [0.8 0.8 0.8]);

gbh = @() eval('grid on; box on; hold on');

runsearch = 0;
if not(runsearch)
    load('CP_2026_02_18_17_33_39.mat')
else
    % Start computation time record
    tic
    %% Initialisation

    % Number of harmonic terms:
    m = 4;

    % Natural frequency:
    omega_0 = 1;

    % Interested damping range
    zeta_array = 0.02:0.02:1;


    %% Particles random generation
    num_samples = 100;
    samples(:,1) = 1*ones(num_samples,1);
    samples(:,2:m) = 0.1*rand(num_samples,m-1);
    samples(1,1:m) = zeros(1,m);
    samples(1,1) = 1;

    % Initialise harmonic timeseries
    N = 501;
    tnorm = linspace(0,1,N);
    [xnorm, xdnorm, xddnorm] = harms(m,N);

    % Create optimisation variable
    a = optimvar('a',m); % m-by-1 variable

    % Tolerances
    general_tolerance = 1e-7;
    weaker_tolerance = 1e-6;

    % Optimisation settings:
    options_sqp = optimoptions(...
        @fmincon, ...
        'ConstraintTolerance', general_tolerance, ...
        'OptimalityTolerance', general_tolerance, ...
        'StepTolerance', general_tolerance, ...
        'MaxFunctionEvaluations', 500, ...
        'Algorithm', 'sqp', ...
        'Display','none');
    options_act = optimoptions(...
        @fmincon, ...
        'ConstraintTolerance', general_tolerance, ...
        'OptimalityTolerance', general_tolerance, ...
        'StepTolerance', general_tolerance, ...
        'MaxFunctionEvaluations', 500, ...
        'Algorithm', 'active-set', ...
        'Display','none');

    %% First continuation loop: upwards
    % Upwards frequency array:
    omega_array_up = 1:0.02:3;
    omega_array = omega_array_up;

    % Data storage arrays:
    std_zero_store = zeros(length(omega_array), length(zeta_array));

    cont_sqp_up.success_mat    = std_zero_store;
    cont_sqp_up.amplitude_mat  = std_zero_store;
    cont_sqp_up.p_res_mat      = std_zero_store;
    cont_sqp_up.a_mat          = zeros(length(omega_array), length(zeta_array), m);


    % Loop:
    for i_zeta = 1:length(zeta_array)
        zeta = zeta_array(i_zeta);

        start_flag = 1;

        for i_omega = 1:length(omega_array)
            iter = 1;
            omega = omega_array(i_omega);

            % Print
            fprintf(['upwards: zeta ' num2str(zeta) ' omega ' num2str(omega) '\n'])

            % Create optimisation problem:
            % Convert main evaulation function into optimisation expressions:
            [min_p,max_p,p,f,x,minx,maxx,F,G] = fcn2optimexpr(@power, zeta, omega_0, omega, a, xnorm, xdnorm, xddnorm, m, N);

            % Define Objective Functions
            prob = optimproblem;
            prob.Objective =  - min_p / max_p; % question

            % Define constraints
            prob.Constraints.normalisation = maxx - minx == 2;
            prob.Constraints.a1 = a(1) * a(1) >= 0.25;

            %% two optimisation methods
            % Optimisation solution SQP:
            % Initial guess:
            if start_flag == 1
                x0.a = zeros(1,m);
                x0.a(1) = 1;
                start_flag = 0;
            else
                x0.a = sol_sqp_up.a;
            end
            [sol_sqp_up,~,~,~] = solve(prob, x0, 'Options', options_sqp);

            % Evaluate all properties of solution:
            [...
                sol_sqp_up.min_p, ...
                sol_sqp_up.max_p, ...
                sol_sqp_up.p, ...
                sol_sqp_up.f, ...
                sol_sqp_up.x, ...
                sol_sqp_up.minx, ...
                sol_sqp_up.maxx, ~, ~] = power(zeta, omega_0, omega, sol_sqp_up.a, xnorm, xdnorm, xddnorm, m, N);

            % Check if solved:
            if  abs(sol_sqp_up.min_p / sol_sqp_up.max_p) <= weaker_tolerance
                % Store data:
                cont_sqp_up.success_mat(i_omega, i_zeta)    = 1;
            elseif iter == 1
                % try all particles
                for iter = 1 : num_samples
                    % Initial guess:
                    x0.a = samples(iter,:);

                    % Optimisation solution, active set method:
                    [sol_sqp_up,~,~,~] = solve(prob, x0, 'Options', options_sqp);

                    % Evaluate all properties of solution:
                    [...
                        sol_sqp_up.min_p, ...
                        sol_sqp_up.max_p, ...
                        sol_sqp_up.p, ...
                        sol_sqp_up.f, ...
                        sol_sqp_up.x, ...
                        sol_sqp_up.minx, ...
                        sol_sqp_up.maxx, ~, ~] = power(zeta, omega_0, omega, sol_sqp_up.a, xnorm, xdnorm, xddnorm, m, N);

                    % Check if solved:
                    if  abs(sol_sqp_up.min_p / sol_sqp_up.max_p) <= weaker_tolerance
                        % Store data:
                        cont_sqp_up.success_mat(i_omega, i_zeta)    = 1;
                        break;
                    end
                end

                %% Only try act optimization if sqp could not find a solution
                if iter == num_samples && abs(sol_sqp_up.min_p / sol_sqp_up.max_p) > weaker_tolerance
                    i_omega_exin = i_omega;
                    for i_omega_ex = i_omega_exin:length(omega_array)
                        x0.a = zeros(1,m);
                        x0.a(1) = 1;
                        omega = omega_array(i_omega_ex);

                        [min_p,max_p,p,f,x,minx,maxx,F,G] = fcn2optimexpr(@power, zeta, omega_0, omega, a, xnorm, xdnorm, xddnorm, m, N);

                        % Define Objective Functions
                        prob = optimproblem;
                        prob.Objective =  - min_p / max_p; % question

                        % Define constraints
                        prob.Constraints.normalisation = maxx-minx == 2;
                        % prob.Constraints.normalisation = a(1)^2 + a(2)^2 + a(3)^2 + a(4)^2 == 1;
                        prob.Constraints.a1 = a(1) * a(1) >= 0.25;

                        % Optimisation solution, sqp method:
                        % -------------------------------------------------------------------------
                        [sol_sqp_up,~,~,~] = solve(prob, x0, 'Options', options_sqp);
                        % Evaluate all properties of solution:
                        [...
                            sol_sqp_up.min_p, ...
                            sol_sqp_up.max_p, ...
                            sol_sqp_up.p, ...
                            sol_sqp_up.f, ...
                            sol_sqp_up.x, ...
                            sol_sqp_up.minx, ...
                            sol_sqp_up.maxx, ~, ~] = power(zeta, omega_0, omega, sol_sqp_up.a, xnorm, xdnorm, xddnorm, m, N);

                        % Optimisation solution, active set method:
                        % -------------------------------------------------------------------------
                        [sol_act_up,~,~,~] = solve(prob, x0, 'Options', options_act);
                        % Evaluate all properties of solution:
                        [...
                            sol_act_up.min_p, ...
                            sol_act_up.max_p, ...
                            sol_act_up.p, ...
                            sol_act_up.f, ...
                            sol_act_up.x, ...
                            sol_act_up.minx, ...
                            sol_act_up.maxx, ~, ~] = power(zeta, omega_0, omega, sol_act_up.a, xnorm, xdnorm, xddnorm, m, N);

                    % if act can find a better solution, then give this solution to sqp
                    if abs(sol_sqp_up.min_p / sol_sqp_up.max_p) > abs(sol_act_up.min_p / sol_act_up.max_p)
                            sol_sqp_up = sol_act_up;
                    end
                    
                    cont_sqp_up.p_res_mat(i_omega_ex, i_zeta)      = abs(sol_sqp_up.min_p / sol_sqp_up.max_p);
                    cont_sqp_up.amplitude_mat(i_omega_ex, i_zeta)  = sol_sqp_up.maxx - sol_sqp_up.minx;
                    cont_sqp_up.a_mat(i_omega_ex, i_zeta, :)       = sol_sqp_up.a;
                    end

                    break;
                end
            end
            cont_sqp_up.p_res_mat(i_omega, i_zeta)      = abs(sol_sqp_up.min_p / sol_sqp_up.max_p);
            cont_sqp_up.amplitude_mat(i_omega, i_zeta)  = sol_sqp_up.maxx - sol_sqp_up.minx;
            cont_sqp_up.a_mat(i_omega, i_zeta, :)       = sol_sqp_up.a;
        end
    end


    %% Second continuation loop: downwards
    % Upwards frequency array:
    omega_array_down = 1:-0.02:0.02;
    omega_array = omega_array_down;

    % Data storage arrays:
    std_zero_store = zeros(length(omega_array), length(zeta_array));

    cont_sqp_down.success_mat    = std_zero_store;
    cont_sqp_down.amplitude_mat  = std_zero_store;
    cont_sqp_down.p_res_mat      = std_zero_store;
    cont_sqp_down.a_mat          = zeros(length(omega_array), length(zeta_array), m);

    % Loop:
    for i_zeta = 1:length(zeta_array)
        zeta = zeta_array(i_zeta);

        start_flag = 1;

        for i_omega = 1:length(omega_array)
            iter = 1;
            omega = omega_array(i_omega);

            % Print
            fprintf(['downwards: zeta ' num2str(zeta) ' omega ' num2str(omega) '\n'])

            % Create optimisation problem:
            % Convert main evaulation function into optimisation expressions:
            [min_p,max_p,p,f,x,minx,maxx,F,G] = fcn2optimexpr(@power, zeta, omega_0, omega, a, xnorm, xdnorm, xddnorm, m, N);

            % Define Objective Functions
            prob = optimproblem;
            prob.Objective =  - min_p / max_p; % question

            % Define constraints
            prob.Constraints.normalisation = maxx-minx == 2;
            prob.Constraints.a1 = a(1) * a(1) >= 0.25;

            %% two optimisation methods
            % Optimisation solution SQP:
            % Initial guess:
            if start_flag ==1
                x0.a = zeros(1,m);
                x0.a(1) = 1;
                start_flag = 0;
            else
                x0.a = sol_sqp_down.a;
            end


            [sol_sqp_down,~,~,~] = solve(prob, x0, 'Options', options_sqp);

            % Evaluate all properties of solution:
            [...
                sol_sqp_down.min_p, ...
                sol_sqp_down.max_p, ...
                sol_sqp_down.p, ...
                sol_sqp_down.f, ...
                sol_sqp_down.x, ...
                sol_sqp_down.minx, ...
                sol_sqp_down.maxx, ~, ~] = power(zeta, omega_0, omega, sol_sqp_down.a, xnorm, xdnorm, xddnorm, m, N);

            % Check if solved:
            if  abs(sol_sqp_down.min_p / sol_sqp_down.max_p) <= weaker_tolerance
                % Store data:
                cont_sqp_down.success_mat(i_omega, i_zeta)    = 1;
            elseif iter == 1
                for iter = 1:num_samples
                    % Initial guess:
                    x0.a = samples(iter,:);

                    % Optimisation solution, active set method:
                    [sol_sqp_down,~,~,~] = solve(prob, x0, 'Options', options_sqp);

                    % Evaluate all properties of solution:
                    [...
                        sol_sqp_down.min_p, ...
                        sol_sqp_down.max_p, ...
                        sol_sqp_down.p, ...
                        sol_sqp_down.f, ...
                        sol_sqp_down.x, ...
                        sol_sqp_down.minx, ...
                        sol_sqp_down.maxx, ~, ~] = power(zeta, omega_0, omega, sol_sqp_down.a, xnorm, xdnorm, xddnorm, m, N);

                    % Check if solved:
                    if  abs(sol_sqp_down.min_p / sol_sqp_down.max_p) <= weaker_tolerance
                        % Store data:
                        cont_sqp_down.success_mat(i_omega, i_zeta)    = 1;
                        break;
                    end
                end

                %% Only try act optimization if sqp could not find a solution
                if iter == num_samples && abs(sol_sqp_down.min_p / sol_sqp_down.max_p) > weaker_tolerance
                    i_omega_exin = i_omega;
                    for i_omega_ex = i_omega_exin:length(omega_array)
                        x0.a = zeros(1,m);
                        x0.a(1) = 1;
                        omega = omega_array(i_omega_ex);

                        [min_p,max_p,p,f,x,minx,maxx,F,G] = fcn2optimexpr(@power, zeta, omega_0, omega, a, xnorm, xdnorm, xddnorm, m, N);

                        % Define Objective Functions
                        prob = optimproblem;
                        prob.Objective =  - min_p / max_p; % question

                        % Define constraints
                        prob.Constraints.normalisation = maxx-minx == 2;
                        % prob.Constraints.normalisation = a(1)^2 + a(2)^2 + a(3)^2 + a(4)^2 == 1;
                        prob.Constraints.a1 = a(1) * a(1) >= 0.25;

                        %% Optimisation solution, sqp method:
                        % -------------------------------------------------------------------------
                        [sol_sqp_down,~,~,~] = solve(prob, x0, 'Options', options_sqp);
                        % Evaluate all properties of solution:
                        [...
                            sol_sqp_down.min_p, ...
                            sol_sqp_down.max_p, ...
                            sol_sqp_down.p, ...
                            sol_sqp_down.f, ...
                            sol_sqp_down.x, ...
                            sol_sqp_down.minx, ...
                            sol_sqp_down.maxx, ~, ~] = power(zeta, omega_0, omega, sol_sqp_down.a, xnorm, xdnorm, xddnorm, m, N);

                        %% Optimisation solution, act method:
                        % -------------------------------------------------------------------------
                        [sol_act_down,~,~,~] = solve(prob, x0, 'Options', options_act);
                        % Evaluate all properties of solution:
                        [...
                            sol_act_down.min_p, ...
                            sol_act_down.max_p, ...
                            sol_act_down.p, ...
                            sol_act_down.f, ...
                            sol_act_down.x, ...
                            sol_act_down.minx, ...
                            sol_act_down.maxx, ~, ~] = power(zeta, omega_0, omega, sol_act_down.a, xnorm, xdnorm, xddnorm, m, N);

                         % if act can find a better solution, then give this solution to sqp
                         if abs(sol_sqp_down.min_p / sol_sqp_down.max_p) > abs(sol_act_down.min_p / sol_act_down.max_p)
                            sol_sqp_down = sol_act_down;
                         end
                        cont_sqp_down.p_res_mat(i_omega_ex, i_zeta)      = abs(sol_sqp_down.min_p / sol_sqp_down.max_p);
                        cont_sqp_down.amplitude_mat(i_omega_ex, i_zeta)  = sol_sqp_down.maxx - sol_sqp_down.minx;
                        cont_sqp_down.a_mat(i_omega_ex, i_zeta, :)       = sol_sqp_down.a;
                    end

                    break;
                end
            end
            cont_sqp_down.p_res_mat(i_omega, i_zeta)      = abs(sol_sqp_down.min_p / sol_sqp_down.max_p);
            cont_sqp_down.amplitude_mat(i_omega, i_zeta)  = sol_sqp_down.maxx - sol_sqp_down.minx;
            cont_sqp_down.a_mat(i_omega, i_zeta, :)       = sol_sqp_down.a;
        end
    end


    %% Merge upward and downward continuation
    % Combined omega array:
    omega_array_comb = [fliplr(omega_array_up), omega_array_down(2:end)];
    Lcomb = length(omega_array_comb);
    cont_comb.success_mat = [flipud(cont_sqp_up.success_mat); cont_sqp_down.success_mat(2:end,:)];
    cont_comb.p_res_mat = [flipud(cont_sqp_up.p_res_mat); cont_sqp_down.p_res_mat(2:end,:)];
    cont_comb.amplitude_mat = [flipud(cont_sqp_up.amplitude_mat); cont_sqp_down.amplitude_mat(2:end,:)];
    cont_comb.a_mat = [flipud(cont_sqp_up.a_mat); cont_sqp_down.a_mat(2:end,:,:)];

    % Stop computation time record
    elapsedTime = toc;
    % Save in timestamped data file:
    save(['CP_' sprintf('%s', datetime('now', 'Format', 'yyyy_MM_dd_HH_mm_ss')) '.mat'])
end

numOnes  = sum(cont_comb.success_mat(:) == 1);
fprintf(['the number of feasible points  '  num2str(numOnes) '\n'])

%% End of escape clause: now plot results
figure(1);
hold on; box on;
colormap([1, 1, 1; [255 0 255]/255]);       % 0 → white, 1 → blue
image(zeta_array, omega_array_comb, cont_comb.success_mat, 'CDataMapping', 'scaled');
set(gca,'YDir','normal'),
title('Composite: Active-set + SQP')
ylabel('Frequency modulation, $\Omega/\Omega_0$')
xlabel('Damping ratio, $\zeta$')


function [min_p,max_p,p,f,x,minx,maxx,F,G] = power(zeta, omega_0, omega, a, xnorm, xdnorm, xddnorm, m, N)

x = zeros(1,N);
dx = zeros(1,N);
ddx = zeros(1,N);

for aindex = 1:m
    x = x + a(aindex)               * xnorm(aindex,:);
    dx = dx + a(aindex)   * omega   * xdnorm(aindex,:);
    ddx = ddx + a(aindex) * omega^2 * xddnorm(aindex,:);
end

% dynamic model
f = ddx + 2 * zeta * omega_0 * dx + omega_0^2 * x;
p = f .* dx;
min_p = min(p);
max_p = max(p);

G = ddx + 2 * zeta * omega_0 * dx;
F = -omega_0^2 * x;

%added Equality Condition
maxx = max(x);
minx = min(x);
end

function [xnorm, xdnorm, xddnorm] = harms(m,N)

xnorm = zeros(m,N);
xdnorm = zeros(m,N);
xddnorm = zeros(m,N);

wt = linspace(0, 2*pi, N);

for index_term = 1:m
    xnorm(index_term,:) =                              cos((2 * index_term - 1) * wt);
    xdnorm(index_term,:) =  - (2 * index_term - 1)   * sin((2 * index_term - 1) * wt);
    xddnorm(index_term,:) = - (2 * index_term - 1)^2 * cos((2 * index_term - 1) * wt);
end
end
```
