## Supplementary material for "Distributed elasticity: a high-reward, moderate-risk strategy for efficient control modulation in insect flight": Code for the work: particle_swarm.m

```
%%%%%%%%%%%%%%%%%%%%%%%%%%%%%%%%%%%%%%%%%
% Particle swarm method for Mapping the space of band-type resonance in insect and insect-inspired
% f = DDX + 2 * zeta * omega_0 * DX + omega_0^2 * X;
% p = f .* DX;
% We aim to find states with p>0
% Author: Liang Wang, Congxiao Zhang, Najmeh Asadimoghaddam, and Arion Pons
% Date: 18th February 2026
% -------------------------------------------------------------------------
clear all
close all
clc;

%% Graphics
% Colours, neon colourset:
NE_pi = [217 12  220]/255;
NE_cy = [4   236 239]/255;
NE_na = [75  88  167]/255;
NE_bk = [0   0   0  ]/255;
NE_wh = [1   1   1  ];
NE_gr = 0.3*[1 1 1];
NE_gm = 0.5*[1 1 1];
NE_gl = 0.7*[1 1 1];

NE_pibk = 0.7*NE_pi + 0.3*NE_bk;
NE_cybk = 0.7*NE_cy + 0.3*NE_bk;
NE_piwh = 0.4*NE_pi + 0.6*NE_wh;
NE_cywh = 0.4*NE_cy + 0.6*NE_wh;
NE_nawh = 0.6*NE_na + 0.4*NE_wh;

% Defaults:
set(groot, 'defaultAxesTickLabelInterpreter','latex');
set(groot, 'defaultLegendInterpreter','latex');
set(groot, 'defaultTextInterpreter','latex');
set(groot, 'defaultTextFontSize', 11);
set(groot, 'defaultAxesXColor', [0 0 0]);
set(groot, 'defaultAxesYColor', [0 0 0]);
set(groot, 'defaultAxesGridAlpha', 1);
set(groot, 'defaultAxesMinorGridAlpha', 1);
set(groot, 'defaultAxesGridColor', [0.8 0.8 0.8]);

gbh = @() eval('grid on; box on; hold on');

%% Select analysis type

% Choose whether to run search or load datafile:
runsearch = 0;
if not(runsearch)
    load('PSO_2025_09_08_21_34_10.mat')
    % load('PSO_2025_08_25_15_29_12.mat')
else
    %Start computation time record  
    tic

    %% Initialisation

    % Number of harmonic terms:
    m = 4;

    % Natural frequency:
    omega_0 = 1;

    % Interested damping range
    zeta_array = 0.02:0.02:1;
    % Interested frequency range
    omega_array = 2:-0.02:0.02;

    % Data storage arrays:
    std_zero_store = zeros(length(omega_array), length(zeta_array));

    act.success_mat    = std_zero_store;
    act.amplitude_mat  = std_zero_store;
    act.p_res_mat      = std_zero_store;

    sqp.success_mat    = std_zero_store;
    sqp.amplitude_mat  = std_zero_store;
    sqp.p_res_mat      = std_zero_store;

    act.a_mat = zeros(length(omega_array), length(zeta_array), m);
    sqp.a_mat = zeros(length(omega_array), length(zeta_array), m);

    comb.success_mat   = std_zero_store;
    comb.p_res_mat     = std_zero_store;
    comb.amplitude_mat = std_zero_store;
    comb.a_mat         = zeros(length(omega_array), length(zeta_array), m);

    %% particles random generation
    num_samples = 2;
    samples(:,1) = 1*ones(num_samples,1);
    samples(:,2:m) = 0.1*rand(num_samples,m-1);
    samples(1,1:m) = zeros(1,m);
    samples(1,1) = 1;

    %% First loop analysis

    % Initialise harmonic timeseries
    N = 501;
    tnorm = linspace(0,1,N);
    [xnorm, xdnorm, xddnorm] = harms(m,N);

    % Create optimisation variable
    a = optimvar('a',m); % m-by-1 variable

    % Index for table:
    table_index = 1;

    % Loop
    for i_zeta = 1:length(zeta_array)
        zeta = zeta_array(i_zeta);

        for i_omega = 1:length(omega_array)
            omega = omega_array(i_omega);

            % Create optimisation problem:
            % Convert main evaulation function into optimisation expressions:
            [min_p,max_p,p,f,x,minx,maxx,F,G] = fcn2optimexpr(@power, zeta, omega_0, omega, a, xnorm, xdnorm, xddnorm, m, N);

            % Define Objective Functions
            prob = optimproblem;
            prob.Objective =  - min_p / max_p; % question

            % Define constraints
            prob.Constraints.normalisation = maxx-minx == 2;
            prob.Constraints.a1 = a(1) * a(1) >= 0.25;

            % Tolerances
            general_tolerance = 1e-7;
            weaker_tolerance = 1e-6;

            % Optimisation settings:
            options_sqp = optimoptions(...
                @fmincon, ...
                'ConstraintTolerance', general_tolerance, ...
                'OptimalityTolerance', general_tolerance, ...
                'StepTolerance', general_tolerance, ...
                'MaxFunctionEvaluations', 500, ...
                'Algorithm', 'sqp', ...
                'Display','none');

            options_act = optimoptions(...
                @fmincon, ...
                'ConstraintTolerance', general_tolerance, ...
                'OptimalityTolerance', general_tolerance, ...
                'StepTolerance', general_tolerance, ...
                'MaxFunctionEvaluations', 500, ...
                'Algorithm', 'active-set', ...
                'Display','none');

            %% Gaussian sampling loop - for the active set method:
            % -------------------------------------------------------------------------
            for iter = 1:num_samples

                % Initial guess:
                x0.a = samples(iter,:);

                % Optimisation solution, active set method:
                [sol_act, ~ , ~, ~] = solve(prob, x0, 'Options', options_act);

                % Evaluate all properties of solution:
                [...
                    sol_act.min_p, ...
                    sol_act.max_p, ...
                    sol_act.p, ...
                    sol_act.f, ...
                    sol_act.x, ...
                    sol_act.minx, ...
                    sol_act.maxx, ~, ~] = power(zeta, omega_0, omega, sol_act.a, xnorm, xdnorm, xddnorm, m, N);

                % Store solution:
                act.p_res_mat(i_omega, i_zeta)      = abs(sol_act.min_p / sol_act.max_p);
                act.amplitude_mat(i_omega, i_zeta)  = sol_act.maxx - sol_act.minx;
                act.a_mat(i_omega, i_zeta, :)       = sol_act.a;

                % Check if solved, and then flag and break:
                if  abs(sol_act.min_p / sol_act.max_p) <= weaker_tolerance
                    % Flag:
                    act.success_mat(i_omega, i_zeta)    = 1;

                    % Break gaussian sampling loop:
                    break;

                end
            end

            %% Gaussian sampling loop - for the SQP method:
            % -------------------------------------------------------------------------
            for iter = 1:num_samples

                % Initial guess:
                x0.a = samples(iter,:);

                % Optimisation solution, SQP method:
                [sol_sqp, ~ , ~, ~] = solve(prob, x0, 'Options', options_sqp);

                % Evaluate all properties of solution:
                [...
                    sol_sqp.min_p, ...
                    sol_sqp.max_p, ...
                    sol_sqp.p, ...
                    sol_sqp.f, ...
                    sol_sqp.x, ...
                    sol_sqp.minx, ...
                    sol_sqp.maxx, ~, ~] = power(zeta, omega_0, omega, sol_sqp.a, xnorm, xdnorm, xddnorm, m, N);

                % Store solution:
                sqp.p_res_mat(i_omega, i_zeta)      = abs(sol_sqp.min_p / sol_sqp.max_p);
                % sqp.p_res_mat(i_omega, i_zeta)      = sol_sqp.min_p / sol_sqp.max_p;
                sqp.amplitude_mat(i_omega, i_zeta)  = sol_sqp.maxx - sol_sqp.minx;
                sqp.a_mat(i_omega, i_zeta, :)       = sol_sqp.a;

                % Check if solved, and then flag and break:
                if  abs(sol_sqp.min_p / sol_sqp.max_p) <= weaker_tolerance
                    % Flag:
                    sqp.success_mat(i_omega, i_zeta)     = 1;
                    % Break gaussian sampling loop:
                    break;

                end
            end

            %% Combine SQP and ACT solutions
            % -------------------------------------------------------------------------
            % Check SQP success first:
            if sqp.success_mat(i_omega, i_zeta) == 1

                comb.success_mat(i_omega, i_zeta)    = 1;
                comb.p_res_mat(i_omega, i_zeta)      = sqp.p_res_mat(i_omega, i_zeta);
                comb.amplitude_mat(i_omega, i_zeta)  = sqp.amplitude_mat(i_omega, i_zeta);
                comb.a_mat(i_omega, i_zeta, :)       = sqp.a_mat(i_omega, i_zeta, :);

                % Check active set success next:
            elseif act.success_mat(i_omega, i_zeta) == 1

                comb.success_mat(i_omega, i_zeta)    = 1;
                comb.p_res_mat(i_omega, i_zeta)      = act.p_res_mat(i_omega, i_zeta);
                comb.amplitude_mat(i_omega, i_zeta)  = act.amplitude_mat(i_omega, i_zeta);
                comb.a_mat(i_omega, i_zeta, :)       = act.a_mat(i_omega, i_zeta, :);

                % If unsucessful, but active set is better:
            elseif act.p_res_mat(i_omega, i_zeta) < sqp.p_res_mat(i_omega, i_zeta)

                comb.p_res_mat(i_omega, i_zeta)      = act.p_res_mat(i_omega, i_zeta);
                comb.amplitude_mat(i_omega, i_zeta)  = act.amplitude_mat(i_omega, i_zeta);
                comb.a_mat(i_omega, i_zeta, :)       = act.a_mat(i_omega, i_zeta, :);

                % If otherwise unsucessful, use SQP:
            else
                comb.p_res_mat(i_omega, i_zeta)      = sqp.p_res_mat(i_omega, i_zeta);
                comb.amplitude_mat(i_omega, i_zeta)  = sqp.amplitude_mat(i_omega, i_zeta);
                comb.a_mat(i_omega, i_zeta, :)       = sqp.a_mat(i_omega, i_zeta, :);

            end

            fprintf(['initial search, ' num2str(i_zeta) '/' num2str(length(zeta_array)) '-' num2str(i_omega) ' complete \n'])
        end
    end


    % Stop computation time record
    elapsedTime = toc;
    % Save in timestamped data file:
    save(['PSO_' sprintf('%s', datetime('now', 'Format', 'yyyy_MM_dd_HH_mm_ss')) '.mat'])

end

%% End of escape clause: now plot results
numOnes  = sum(comb.success_mat == 1);
fprintf(['the number of feasible points  '  num2str(numOnes) '\n'])

% Map of optimised states across different methods:
% incl. band-type resonant states, and others:
figure(1);
hold on; box on;
colormap([1, 1, 1; [255 0 255]/255]);       % 0 → white, 1 → blue
image(zeta_array, omega_array, comb.success_mat, 'CDataMapping', 'scaled');
set(gca,'YDir','normal'),
title('Particle Swarm: Active-set + SQP')
ylabel('Frequency modulation, $\Omega/\Omega_0$')
xlabel('Damping ratio, $\zeta$')


%% Functions

function [min_p,max_p,p,f,x,minx,maxx,F,G] = power(zeta, omega_0, omega, a, xnorm, xdnorm, xddnorm, m, N)

x = zeros(1,N);
dx = zeros(1,N);
ddx = zeros(1,N);

for aindex = 1:m
    x = x + a(aindex)               * xnorm(aindex,:);
    dx = dx + a(aindex)   * omega   * xdnorm(aindex,:);
    ddx = ddx + a(aindex) * omega^2 * xddnorm(aindex,:);
end

% dynamic model
f = ddx + 2 * zeta * omega_0 * dx + omega_0^2 * x;
% f = ddx + 2 * zeta * omega_0 * abs(dx).* dx + omega_0^2 * x;
p = f .* dx;
min_p = min(p);
max_p = max(p);

G = ddx + 2 * zeta * omega_0 * dx;
F = -omega_0^2 * x;

%added Equality Condition
maxx = max(x);
minx = min(x);
end


function [xnorm, xdnorm, xddnorm] = harms(m,N)

xnorm = zeros(m,N);
xdnorm = zeros(m,N);
xddnorm = zeros(m,N);

wt = linspace(0, 2*pi, N);

for index_term = 1:m
    xnorm(index_term,:) =                              cos((2 * index_term - 1) * wt);
    xdnorm(index_term,:) =  - (2 * index_term - 1)   * sin((2 * index_term - 1) * wt);
    xddnorm(index_term,:) = - (2 * index_term - 1)^2 * cos((2 * index_term - 1) * wt);
end
end
```
