## Supplementary material for "Distributed elasticity: a high-reward, moderate-risk strategy for efficient control modulation in insect flight": Code for the work: readme.txt

The attached three scripts correspond to different numerical methods for mapping the band-type resonance.
'natural_continuation.m' is the script for the natural continuation mapping method.
'particle_swarm.m' is the script for the particle swarm mapping method.
'composite.m' is the script for the composite mapping method.
